# Comparative multi-omics analysis uncovers contrasting molecular profiles of canine and human thyroid carcinomas

**DOI:** 10.1101/2025.03.14.643300

**Authors:** Sunetra Das, Samantha N. Schlemmer, Rupa Idate, Susan E. Lana, Daniel P. Regan, Douglas H. Thamm, Dawn L. Duval

## Abstract

Thyroid tumors represent 1-3% of canine cancers, with most tumors classified as follicular carcinomas and less frequently as medullary carcinomas. In comparison, only 10-15% of human thyroid cancers are follicular and 2% are medullary, with prevalent activating mutations in *BRAF* and *NRAS,* or *RET,* respectively. A cohort of canine thyroid carcinomas underwent histopathological (n=60 and paired molecular (n=30, WES and/or RNAseq) exploration. Clustering of tumor transcriptomes produced 2 groups; T1 and T2 clusters comprised of follicular thyroid carcinomas (FTC) and medullary thyroid carcinomas (MTC), respectively. Tumors were histologically typed as follicular, compact, and follicular-compact on blinded review, with most MTC classified as compact with rare follicular-compact appearance, while FTC displayed all 3 patterns. FTC samples had significantly elevated levels of *ERBB2* and HER2 protein, and *RET* signaling was up-regulated in MTC. Recurrent somatic mutations in *DNMT1*, *STAT2*, *SALL4, HSP90AA1, MEN1, MUC4, THRAP3, CDK4, NOTCH2,* and *THRAP3* were identified in at least 10% of samples. Individual variants were also identified in *KRAS, ARAF, GNAS*, and *ERBB2*. Additionally, we identified fusion genes that included *TG*, *FGFR2*, and *PAX8*. These data suggest that canine MTC, like their human counterparts, may be driven by RET signaling. In contrast, FTC show limited reliance on RAS/RAF signaling, prevalent in human TC, for oncogenic progression. Across the analyzed samples, 60% of tumors had mutations in at least one DNA repair-related pathway, suggesting that the accumulation of DNA damage may drive cancer progression in canine thyroid tumors. Elevated HER2 protein staining was associated with shorter progression free survival (PFS). Thyroid tumor size (>4.25 cm) was also associated with shorter overall survival and PFS, consistent with previous reports; however, metastasis at diagnosis was not correlated with outcome. Additional studies are warranted to explore the utility of these biomarkers to improve diagnosis and treatment of thyroid carcinoma in dogs.

## Introduction

Canine thyroid carcinomas (TC) are endocrine tumors that arise from either thyroid follicular (thyrocytes) or medullary/parafollicular cells (C-cells). The thyrocytes produce thyroglobulin, thyroxine and colloid, and the medullary cells produce calcitonin. Thyroid cancer accounts for 1-4% and 2.2% of all cancer types in dogs and humans, respectively [1], https://seer.cancer.gov/statfacts/html/thyro.html. In both humans and dogs, thyroid cancer is the most common type of endocrine cancer [2]. Thyroid cancer in dogs can be categorized into two types: follicular TC (FTC) which account for approximately 63.6–87.5% of cases, and medullary TC (MTC) which make up about 12.5–36.4% of canine TC [3, 4]. FTCs can be further classified by histologic subtype as follicular (11.9–32.0%), follicular-compact (32.0–58.0%), or compact/solid (13.6–40.5%) based on the World Health Organization (WHO) scheme [4–7]. The compact subtype of FTC can be difficult to distinguish from MTC due to its more solid histologic appearance, necessitating immunohistochemistry, in particular thyroglobulin for FTC and calcitonin or calcitonin-related peptide for MTC [2, 6, 8–10]. Unlike humans, the papillary subtype is infrequently reported in dogs, accounting for approximately 3.7–7.1% of canine TC [7, 10].

The molecular landscapes for several subtypes of human TC have been previously reported [11–15]. In comparison, only a few targeted genetic studies have been reported in dogs [16–18]. The most common cancer drivers in human papillary TC (PTC) are *BRAF* (48%), *NRAS* (7%), and *HRAS* (3%), whereas activating mutations in *RET* or *RAS* drive MTC [11, 19]. Conversely, *RET* fusions have been identified in up to 30% of sporadic PTC and *NTRK* gene fusions in 2-3% of cases [20]. In addition to BRAF mutations, anaplastic TC have recurrent mutations in *TP53* (58%) and *TERT* (54%) [13, 21] . In broad terms, human TC have been categorized as BRAF-like or RAS-like with the BRAF-like tumors exhibiting high levels of MAP kinase pathway activation and typically falling into the papillary subtype, while the RAS-like tumors commonly have a follicular pattern with both MAP kinase and PI3 kinase pathway activation [11]. In addition to the RAS genes, other RAS-like mutational drivers include *BRAF*^K601E^, *DICER1*, *EZH1*, *EIF1AX*, *PTEN* mutations and *PPARG* or *THADA* gene fusions [22]. Although activating mutations in *KRAS* have been identified in 2 out of 43 dogs with TC [16], the remainder of these drivers of human TC have not been identified in canine studies to date.

Currently, the clinical treatment of dogs with TC is not defined by cell origin (i.e., FTC vs MTC). The preferred first-line treatment for canine TC is surgical excision, if feasible, with median survival times of greater than 36 months reported in dogs with non-metastatic, well-encapsulated tumors [23–25]. Radiation therapy (i.e., external beam radiation, radioiodine) and adjunctive chemotherapy and/or immunotherapy have also been used with varying clinical benefit in certain patient cohorts compared to surgery alone [5, 26–28]. Clinical findings that have been associated with worse prognosis include tumor volume greater than 20 cm^3^, bilateral tumors, and presence of tumor vascular invasion. Up to 38% of dogs have metastasis at the time of diagnosis [26], and these dogs may experience shorter survival [29]. The five-year survival rate of human patients with either MTC, FTC, or PTC is relatively good, 90 to 99%, (https://seer.cancer.gov/statfacts/html/thyro.html); however, anaplastic TC is highly aggressive and has a one-year survival rate of 35% [30]. Molecularly targeted agents, including multi-targeted tyrosine kinase inhibitors or RET, TRK, or ALK kinase inhibitors in fusion-positive tumors, have also been utilized in human TC [31]. In contrast, molecular characterization is currently lacking to aid in the diagnosis, prognosis, and targeted treatment of canine TC.

This study documents a comparative analysis of the genomes and transcriptomes of canine and human TC. Pathological exploration of 60 canine TC tumors identified 54 FTC and 6 MTC, of which 30 tumors (25 FTC, 5 MTC) were sequenced via next-generation sequencing technologies. In accordance with previous studies, larger tumor size at diagnosis correlated with shorter progression free intervals (PFI), although many dogs in this cohort demonstrated prolonged survival (median overall survival for all TC >60 months with thyroidectomy alone, with or without metastasis at diagnosis). The mutational profiles from RNAseq and/or WES revealed limited homology to human thyroid tumors. Transcriptomic analysis identified activation of *RET* signaling in MTC and *ERBB2* signaling in FTC tumors, and HER2 gene and protein expression were associated with shorter PFI. The mutational landscape of canine TC shows a heterogenous population of somatic variants and, unlike human TC, lacks recurrently mutated driver genes. The commonality across these tumors lay in uncovering DNA repair pathway genes mutated in 60% of the tumors analyzed. The presence of a heterogeneous mutational landscape, along with *MEN1* somatic variants in 10% of the analyzed samples, parallels the gene variant profiles observed in human neuroendocrine tumors.

### Results and Discussion

#### Clinicopathologic data

Sixty dogs were included in this study, of which survival data were available in 53 cases and advanced sequencing data (WES and/or RNAseq) in 30 cases **(S1 Table).** Clinical and histopathologic data for these animals are summarized in **Tables 1 and 2**, respectively.

**Table 1:**
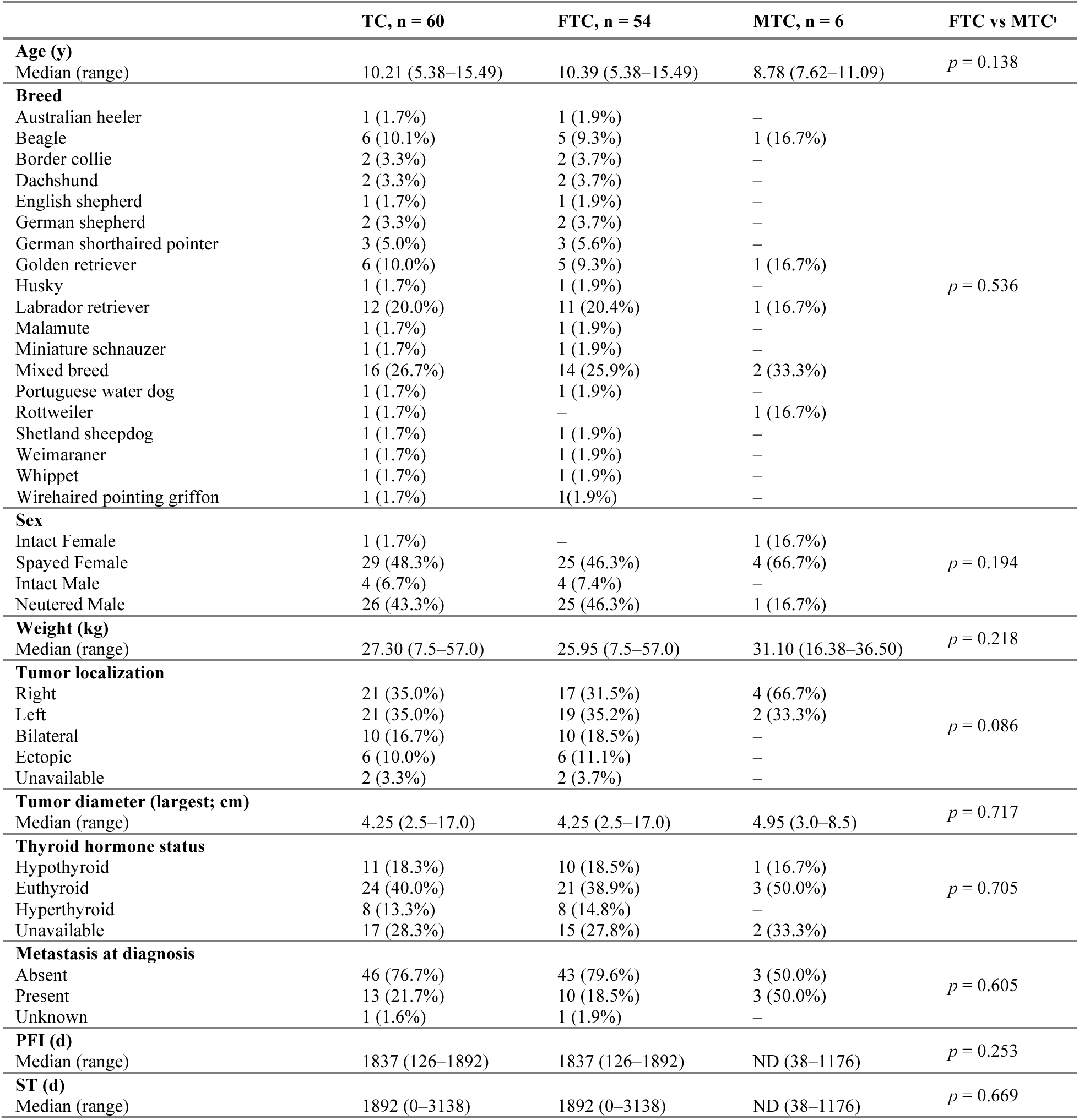
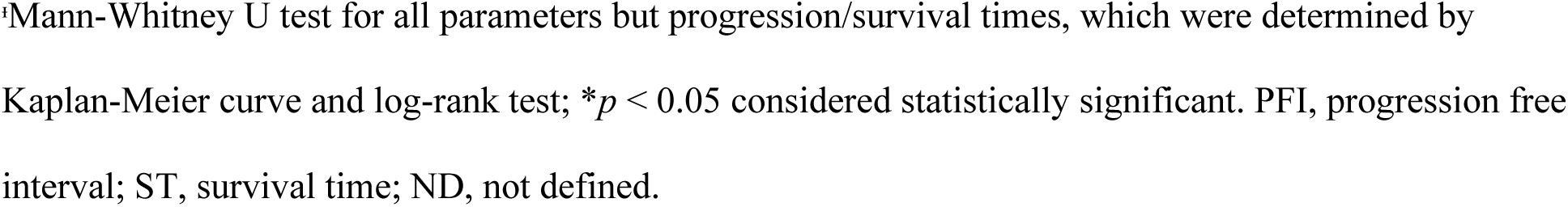
Clinical data from 60 canine thyroid carcinomas (TC), which are also separated into putative follicular and medullary tumors (FTC and MTC, respectively) based on calcitonin expression (see Table 2) for comparisons based on tumor cell origin.

|  | TC, n = 60 | FTC, n = 54 | MTC, n = 6 | FTC vs MTC<sup>a</sup> |
| --- | --- | --- | --- | --- |
| **Age (y)** |  |  |  |  |
| Median (range) | 10.21 (5.38–15.49) | 10.39 (5.38–15.49) | 8.78 (7.62–11.09) | $p = 0.138$ |
| **Breed** |  |  |  |  |
| Australian heeler | 1 (1.7%) | 1 (1.9%) | – | $p = 0.536$ |
| Beagle | 6 (10.1%) | 5 (9.3%) | 1 (16.7%) |  |
| Border collie | 2 (3.3%) | 2 (3.7%) | – |  |
| Dachshund | 2 (3.3%) | 2 (3.7%) | – |  |
| English shepherd | 1 (1.7%) | 1 (1.9%) | – |  |
| German shepherd | 2 (3.3%) | 2 (3.7%) | – |  |
| German shorthaired pointer | 3 (5.0%) | 3 (5.6%) | – |  |
| Golden retriever | 6 (10.0%) | 5 (9.3%) | 1 (16.7%) |  |
| Husky | 1 (1.7%) | 1 (1.9%) | – |  |
| Labrador retriever | 12 (20.0%) | 11 (20.4%) | 1 (16.7%) |  |
| Malamute | 1 (1.7%) | 1 (1.9%) | – |  |
| Miniature schnauzer | 1 (1.7%) | 1 (1.9%) | – |  |
| Mixed breed | 16 (26.7%) | 14 (25.9%) | 2 (33.3%) |  |
| Portuguese water dog | 1 (1.7%) | 1 (1.9%) | – |  |
| Rottweiler | 1 (1.7%) | – | 1 (16.7%) |  |
| Shetland sheepdog | 1 (1.7%) | 1 (1.9%) | – |  |
| Weimaraner | 1 (1.7%) | 1 (1.9%) | – |  |
| Whippet | 1 (1.7%) | 1 (1.9%) | – |  |
| Wirehaired pointing griffon | 1 (1.7%) | 1 (1.9%) | – |  |
| **Sex** |  |  |  |  |
| Intact Female | 1 (1.7%) | – | 1 (16.7%) | $p = 0.194$ |
| Spayed Female | 29 (48.3%) | 25 (46.3%) | 4 (66.7%) |  |
| Intact Male | 4 (6.7%) | 4 (7.4%) | – |  |
| Neutered Male | 26 (43.3%) | 25 (46.3%) | 1 (16.7%) |  |
| **Weight (kg)** |  |  |  |  |
| Median (range) | 27.30 (7.5–57.0) | 25.95 (7.5–57.0) | 31.10 (16.38–36.50) | $p = 0.218$ |
| **Tumor localization** |  |  |  |  |
| Right | 21 (35.0%) | 17 (31.5%) | 4 (66.7%) | $p = 0.086$ |
| Left | 21 (35.0%) | 19 (35.2%) | 2 (33.3%) |  |
| Bilateral | 10 (16.7%) | 10 (18.5%) | – |  |
| Ectopic | 6 (10.0%) | 6 (11.1%) | – |  |
| Unavailable | 2 (3.3%) | 2 (3.7%) | – |  |
| **Tumor diameter (largest; cm)** |  |  |  |  |
| Median (range) | 4.25 (2.5–17.0) | 4.25 (2.5–17.0) | 4.95 (3.0–8.5) | $p = 0.717$ |
| **Thyroid hormone status** |  |  |  |  |
| Hypothyroid | 11 (18.3%) | 10 (18.5%) | 1 (16.7%) | $p = 0.705$ |
| Euthyroid | 24 (40.0%) | 21 (38.9%) | 3 (50.0%) |  |
| Hyperthyroid | 8 (13.3%) | 8 (14.8%) | – |  |
| Unavailable | 17 (28.3%) | 15 (27.8%) | 2 (33.3%) |  |
| **Metastasis at diagnosis** |  |  |  |  |
| Absent | 46 (76.7%) | 43 (79.6%) | 3 (50.0%) | $p = 0.605$ |
| Present | 13 (21.7%) | 10 (18.5%) | 3 (50.0%) |  |
| Unknown | 1 (1.6%) | 1 (1.9%) | – |  |
| **PFI (d)** |  |  |  |  |
| Median (range) | 1837 (126–1892) | 1837 (126–1892) | ND (38–1176) | $p = 0.253$ |
| **ST (d)** |  |  |  |  |
| Median (range) | 1892 (0–3138) | 1892 (0–3138) | ND (38–1176) | $p = 0.669$ |

<sup>a</sup>Mann-Whitney U test for all parameters but progression/survival times, which were determined by Kaplan-Meier curve and log-rank test; \* $p < 0.05$ considered statistically significant. PFI, progression free interval; ST, survival time; ND, not defined.

**Table 2:** Histopathological data from 60 canine thyroid carcinomas (TC), which are also separated into putative follicular and medullary tumors (FTC and MTC, respectively) based on calcitonin expression (IHC and/or transcriptomic) for comparisons based on tumor cell origin.

|  | TC, n = 60 | FTC, n = 54 | MTC, n = 6 | FTC vs MTC<sup>a</sup> |
| --- | --- | --- | --- | --- |
| **Histologic pattern** |  |  |  |  |
| Follicular | 24 (40.0%) | 24 (44.4%) | – | $p = 0.0015^*$ |
| Follicular-compact | 26 (43.3%) | 24 (44.4%) | 2 (33.3%) |  |
| Compact | 10 (16.7%) | 6 (11.1%) | 4 (66.7%) |  |
| **Tumor differentiation** |  |  |  |  |
| Well | 20 (33.3%) | 20 (37.0%) | – | $p = 0.0015^*$ |
| Moderate | 32 (53.3%) | 30 (55.6%) | 2 (33.3%) |  |
| Poor | 8 (13.3%) | 4 (7.4%) | 4 (66.7%) |  |
| **Nuclear atypia** |  |  |  |  |
| Mild | 34 (56.7%) | 33 (61.1%) | 1 (16.7%) | $p = 0.062$ |
| Moderate | 21 (35.0%) | 17 (31.5%) | 4 (66.7%) |  |
| Marked | 5 (8.3%) | 4 (7.4%) | 1 (16.7%) |  |
| **Invasion/infiltration (microscopic)** |  |  |  |  |
| Absent | – | – | – | $p > 0.999$ |
| Present | 60 (100%) | 54 (100%) | 6 (100%) |  |
| **Tumor emboli (microscopic)** |  |  |  |  |
| Absent | 28 (46.7%) | 25 (46.3%) | 3 (50.0%) | $p > 0.999$ |
| Present | 32 (53.3%) | 29 (53.7%) | 3 (50.0%) |  |
| **Necrosis (microscopic)** |  |  |  |  |
| Absent | 21 (35.0%) | 17 (31.5%) | 4 (66.7%) | $p = 0.171$ |
| Present | 39 (65.0%) | 37 (68.5%) | 2 (33.3%) |  |
| **Hemorrhage (microscopic)** |  |  |  |  |
| Absent | 14 (23.0%) | 11 (20.4%) | 2 (33.3%) | $p = 0.602$ |
| Present | 47 (77.0%) | 43 (79.6%) | 4 (66.7%) |  |
| **Mineral/bone (microscopic)** |  |  |  |  |
| Absent | 49 (81.7%) | 44 (81.5%) | 6 (100%) | $p = 0.577$ |
| Present | 11 (18.3%) | 10 (18.5%) | – |  |
| **Calcitonin IHC** |  |  |  |  |
| Negative | 55 (91.7%) | 54 (100%) | 1 (16.7%) | $p < 0.0001^*$ |
| Positive | 5 (8.3%) | – | 5 (83.3%) |  |
| **HER2 IHC Score (ASCO-CAP)[76, 77]** |  |  |  |  |
| 0, negative | 13 (21.7%) | 8 (14.8%) | 5 (83.3%) | $p = 0.0007^*$ |
| 1, negative | 21 (35.0%) | 20 (37.0%) | 1 (16.7%) |  |
| 2, equivocal | 15 (25.0%) | 15 (27.8%) | – |  |
| 3, positive | 11 (18.3%) | 11 (20.4%) | – |  |
| **HER2 IHC Score (Peña)[78]** |  |  |  |  |
| 0, negative | 13 (21.7%) | 8 (14.8%) | 5 (83.3%) | $p = 0.0007^*$ |
| 1, negative | 21 (35.0%) | 20 (37.0%) | 1 (16.7%) |  |
| 2, equivocal | 19 (31.7%) | 19 (35.2%) | – |  |
| 3, positive | 7 (11.7%) | 7 (13.0%) | – |  |

<sup>a</sup>Mann-Whitney U test for all parameters; \* $p < 0.05$ considered statistically significant.

This cohort included 54 (90%) follicular tumors (FTC) and 6 (10%) medullary tumors (MTC) based on calcitonin expression (IHC and/or transcriptomic), similar to previously reported incidence rates [2–4, 8, 9]. Given small tumor specimens, a single marker (calcitonin) was used to determine cell origin on IHC, as previous literature suggested a high concordance of calcitonin immunoreactivity in canine MTC [9, 10]. A single MTC in our cohort had very faint calcitonin immunoreactivity on IHC, which was initially reported as “negative” when reviewed by blinded pathologists but was later determined to be MTC based on high calcitonin transcript expression. The discordant results between IHC and transcript expression in this specimen could be attributed to intratumor heterogeneity as different tissue samples within the tumor were used for IHC and genomic analyses, or less likely rapid release and/or degradation of calcitonin by tumor cells prior to fixation [8]. Calcitonin-negative neuroendocrine TC have been infrequently encountered in the dog [32], and an extended IHC panel to include calcitonin gene related peptide and synaptophysin could have been considered for this specimen but were not pursued due to limited tissue remaining on the FFPE block.

Clinical data (i.e., age at diagnosis, breed, sex, median weight, median tumor size (largest diameter), tumor localization, and thyroid hormone status) were not statistically different between FTC and MTC. Thyroid tumors were predominantly unilateral (70%) without a lobe predilection although no MTC were present in bilateral nor ectopic locations. Approximately 15% of FTC were functional leading to increased serum total T4, a slightly lower incidence than previous reports [9, 27, 33]. No dogs with MTC were documented as hyperthyroid but thyroid hormone testing was unavailable in 2 cases. As previously documented, many of the dogs affected were medium to large breeds over 7 years old, without a sex predilection [27, 34, 35]. The Labrador retriever was the most frequently reported single pure breed in this cohort (n=12), and like previous reports, the beagle and golden retriever were also among commonly affected breeds with FTC [33, 35, 36]. Consistent with the literature, approximately 21% of dogs with TC had metastasis at diagnosis. Although MTC had numerically increased metastasis at diagnosis, the incidence was not significantly different between FTC and MTC. At diagnosis, FTC metastases were found in regional lymph nodes (n=8) and lung (n=2), while all MTC metastases were noted in regional lymph nodes (n=3). The incidence of metastasis and the median tumor size (4.25 cm) at diagnosis likely reflect later detection, as these tumors are often found incidentally, though some animals may show clinical signs such as coughing, dysphagia, or gagging [2, 37].

Tumor histologic pattern (i.e., follicular, follicular-compact, compact arrangement) and tumor differentiation (i.e., well, moderate, poor) were statistically different between FTC and MTC (*p* = 0.0015 for both parameters), with MTC tumors frequently displaying compact/solid pattern and, subjectively, poor differentiation. FTCs were most commonly follicular or follicular-compact (both 44.4%) patterns with rare tumors displaying compact (11.1%) arrangements; a recent review cites follicular-compact as the most frequently encountered histotype [2]. Interestingly, a portion of MTC were classified as follicular-compact pattern on blinded H&E review, possibly due to pseudo-follicle formation and/or entrapment of nonneoplastic follicular cells [8], underscoring the importance of IHC to differentiate FTC from MTC [2, 10]. Other routine histologic assessments, including subjective nuclear atypia (i.e., mild, moderate, marked) and the presence of microscopic tumor emboli, necrosis, hemorrhage, or mineral/bone, were not statistically different between FTC and MTC; although, no MTC contained mineral/bone. Microscopic invasion was noted in 100% and tumor emboli were appreciated histologically in 53% of all TC in this cohort, and many tumors contained hemorrhage and necrosis, which may be related to later diagnosis and larger tumor size.

### RNASeq and Whole Exome Sequencing Analysis

#### Transcriptomic analysis identifies two discrete clusters of canine thyroid carcinoma

The UMAP analysis of 30 canine TC and 5 normal canine thyroid tissues showed two distinct clusters of TC (T1 and T2) and one cluster of normal samples (**Figure 1A**) [38] . Most follicular and follicular-compact (96%) tumors were grouped in the T1 cluster, except for the TC22 sample (compact histological pattern). The other 4 compact tumors were grouped in the T2 cluster as well as a single follicular-compact sample. The T1 or FTC cluster (n=25) included a sample (TC1088) that lacked thyroid-specific transcription factor (*NKX2-1/TTF-1*) (**Figure 1B**). *NKX2-1* is essential for thyroid tissue development and maintenance, and *NKX2-1* loss is correlated with de-differentiated human TC [39], suggesting that TC1088 might represent a de-differentiated follicular carcinoma. The T2 cluster samples were predominantly MTC with high calcitonin levels at the transcript and protein levels (**Figure 1B**, **Table 2**). High levels of serum calcitonin are used as a biomarker for MTC and other thyroid C-cell diseases in humans [40], although a rare pathology of calcitonin-negative MTCs has also been reported [41]. Similarly, hierarchical clustering based only on gene expression of calcitonin A and B and the thyroid transcription factor, *NKX2-1*, grouped the 4 calcitonin-staining tumors (via IHC) along with 3 histologically compact tumors. The remainder of the tumors with moderate to low expression of the 3 markers were clustered together, except for TC1088 which lacked *NKX2-1* expression. Differences in calcitonin staining and expression could be due to tumor heterogeneity or the inclusion of normal parafollicular cells in the tumor sections utilized for RNAseq.

**Fig 1.**
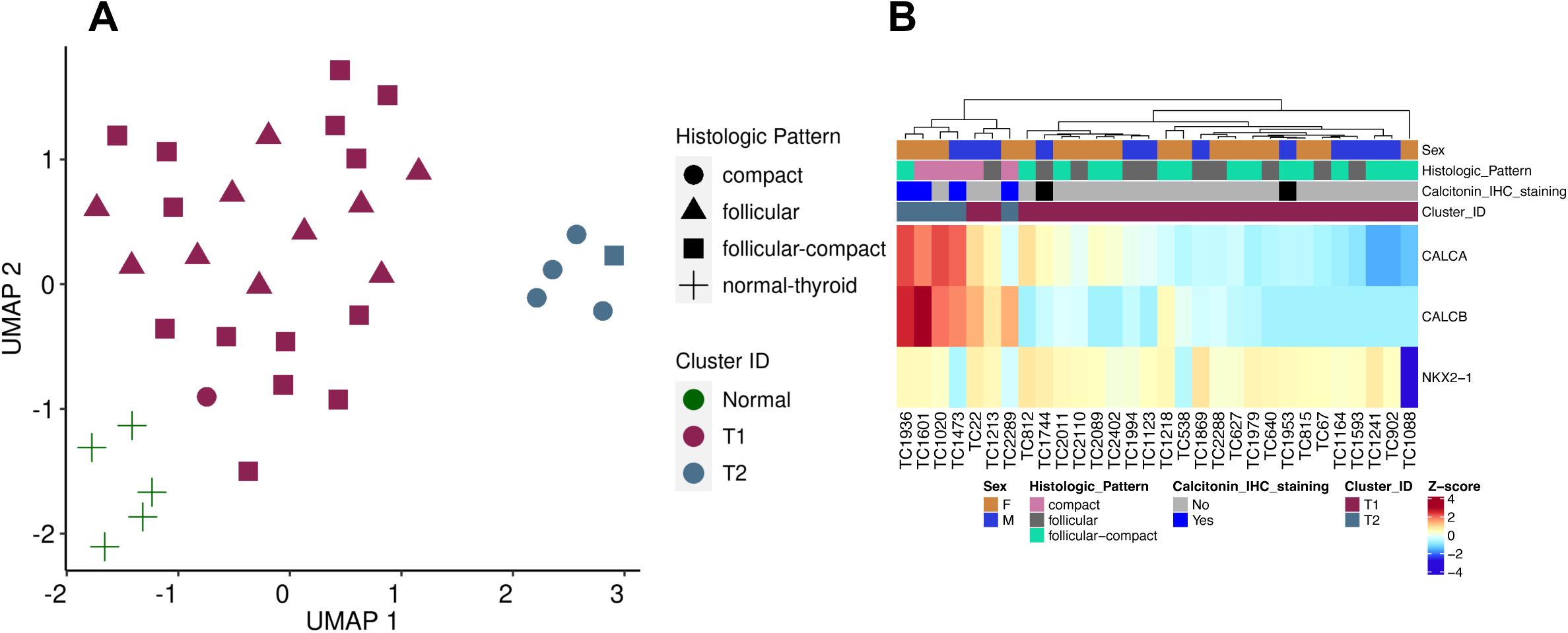
Defining clusters of canine thyroid carcinoma. A. Genes (n=1,013) with highly variant expression levels across thyroid tumor and normal samples were used for UMAP clustering. B. Heatmap of thyroid cell type markers: calcitonin A and B (*CALCA* and *CALCB)* and *NKX2-1* transcripts. The samples with high calcitonin transcript levels also had positive calcitonin IHC staining with exception of TC1020 sample. The loss of thyroid-related transcription factor (NKX2-1) transcript was observed in TC1088 sample.

#### Thyroid differentiation and signaling

We evaluated a Thyroid Differentiation Score (TDS) used in molecular subtyping of human TC, as poorly differentiated papillary tumors are considered to be more aggressive [11]. This score is calculated using expression data of 16 thyroid-specific metabolic and functional genes (**Figure 2A**). Among the canine TC in this study, low TDS was associated with MTCs, and FTCs were associated with higher TDS values. The sample TC1088 (FTC) had very low TDS due to the loss of *NKX2-1*. The association of low to high TDS values with “poor,” “moderate,” and “well” tumor pathological differentiation categories was significant (Kruskal-Wallis, *p* = 0.01, **Figure 2B**).

**Fig 2.**
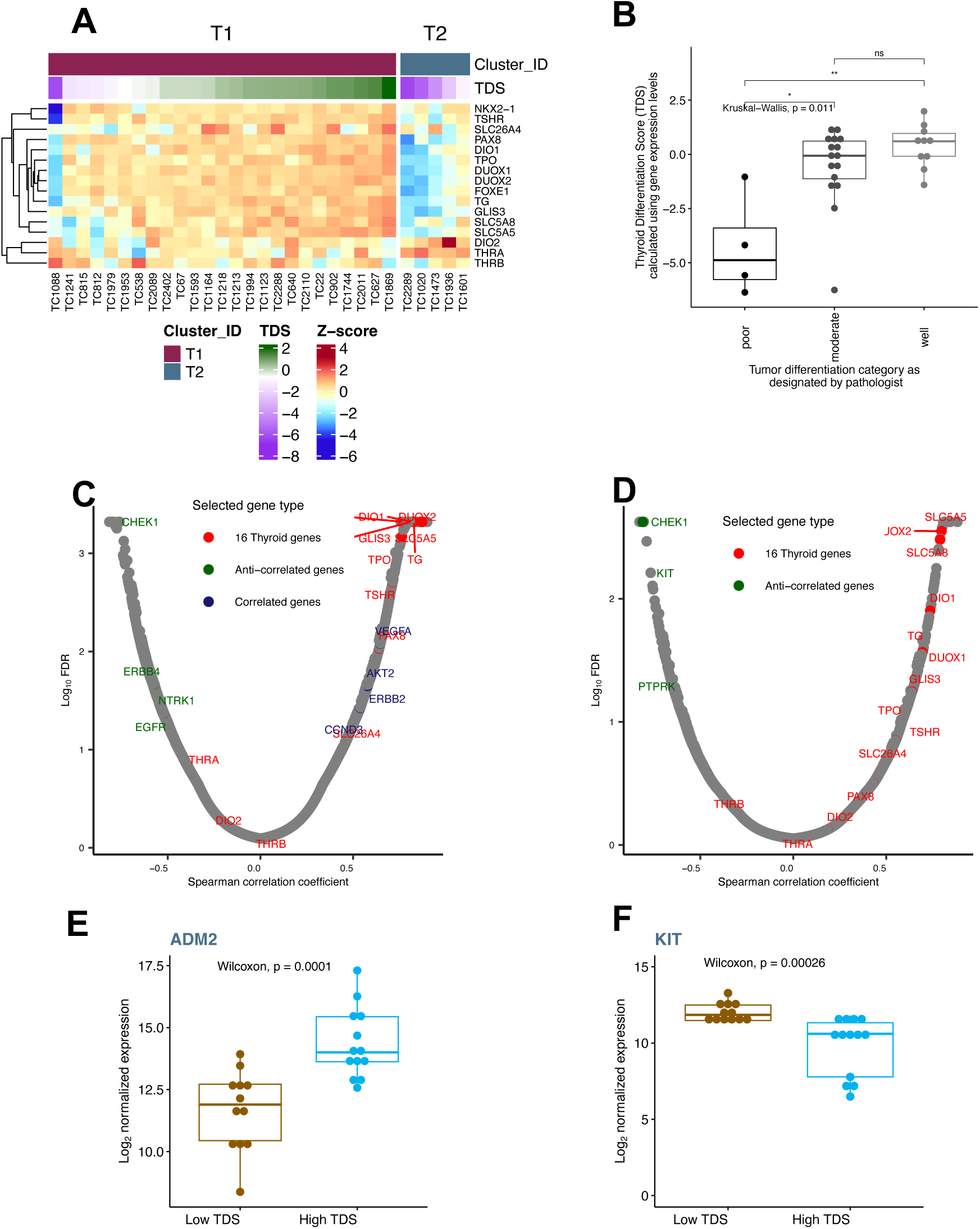
Thyroid differentiation score (TDS) and its association with tumor grade and gene expression. A. Heatmap of 16 thyroid metabolism and function genes used in calculating the TDS. The samples were sorted in ascending order of TDS values. The two groups T1 and T2 represent FTC and MTC tumors, respectively. B. Boxplot showing the association of tumor differentiation grades and TDS. C & D. Transcript expression levels correlated with TDS across all 30 samples (C) and 25 FTC samples (D). The Spearman correlation coefficients were plotted against Log_10_ False Discovery Rate (FDR). The red circles emphasize the 16 thyroid-related genes, and the selected positively- and negatively- or anti-correlated genes are blue and green circles, respectively. E & D. Boxplot of expression data where higher transcript levels were associated with high TDS (ADM2) and low TDS (KIT) within the FTC cohort.

In human tumors, TDS correlates with the *BRAF*^V600E^*-RAS* score (BRS), where low and high TDS were often associated with *BRAF*^V600E^ and RAS mutations in human TC, respectively [11]. In addition, the correlation of expression levels of signaling molecules with TDS in BRAF and RAS cohorts led to the identification of less common mutations in human TC. Similarly, we correlated the global transcriptome with TDS to identify signaling molecules enriched in 30 canine tumors (**Figure 2C**, **S2 Table**). As expected, most of the 16 thyroid-related genes were correlated with TDS, in addition to several cancer-related genes. The key cancer genes positively correlated with TDS were *AKT2*, *VEGFA*, *ERBB2*, and *CCND3.* This reiterates the higher expression levels of *ERBB2* and *AKT2* in FTC samples relative to MTCs. Additionally, TDS anti-correlated genes included *CHEK1*, *ERBB4*, *EGFR,* and *NTRK1* and genes up-regulated in MTC samples. These data suggest that the TDS is specific to tumors deriving from thyroid gland follicular cells.

To identify signaling pathways within FTC (n=25) samples, we repeated the TDS correlation analyses using only the FTC cohort (**Figure 2D, S3 Table**). Although no significantly correlated cancer genes were identified, a tumor angiogenic factor, *ADM2* (Adrenomedullin 2), was associated with a high TDS (well-differentiated) score (**Figure 2E**). ADM2 stimulates PKA (Protein Kinase A) and ERK pathways *in vitro*, promoting cancer cell proliferation and migration. In human TC, *ADM2* is up-regulated in nutrient excess conditions, especially in the context of a high fat diet [42]. Some of the key cancer genes that are anti- or negatively correlated with TDS within the FTC cohort included *KIT* (**Figure 2F**), *CHEK1*, and *PTPRK*. The higher expression of *KIT* in less differentiated canine FTC samples contrasts with *BRAF^V600E^*-driven human PTC samples, where *KIT* expression positively correlates with TDS [11]. As the canine tumors lack homologous *BRAF^V600E^* driver mutations (see results below), the distinct *KIT* expression profile offers valuable insights into alternative mechanisms driving TC in this species compared to humans.

#### Distinct pathways regulate follicular and medullary thyroid carcinomas

We ran a differential expression (DE) analysis between the two clusters of TC. There were 1,348 and 1,050 genes up-regulated in T1/FTC and T2/MTC groups, respectively (**Figure 3A, S4 Table**). Some of the key cancer genes up-regulated in T1/FTC included thyroid gland-specific genes like *TRHR*, *TSHR*, *PAX8*, and fibroblast growth factor receptors, *FGFR2* and *FGFR*. The genes up-regulated in T2/MTC group (i.e., down-regulated in FTC) included *RET*, *NTRK1*, and *MYC* (**Figure 3B, S2 Fig**). Although we did not detect any fusions or short variants in the *RET* gene, the over-expression of *RET* compared to both T1/FTC and normal samples suggests that RET signaling might drive cancer progression in canine MTCs, similar to human MTC (**S2 Fig**). Additionally, these data showed differential expression of the ErbB family of receptor tyrosine kinases in T1/FTC and T2/MTC groups. *ERBB2* was significantly up-regulated in T1/FTC samples, and both *ERBB4* and *EGFR* genes had higher transcript levels in the T2/MTC samples than in normal and T1/FTC groups.

**Fig 3.**
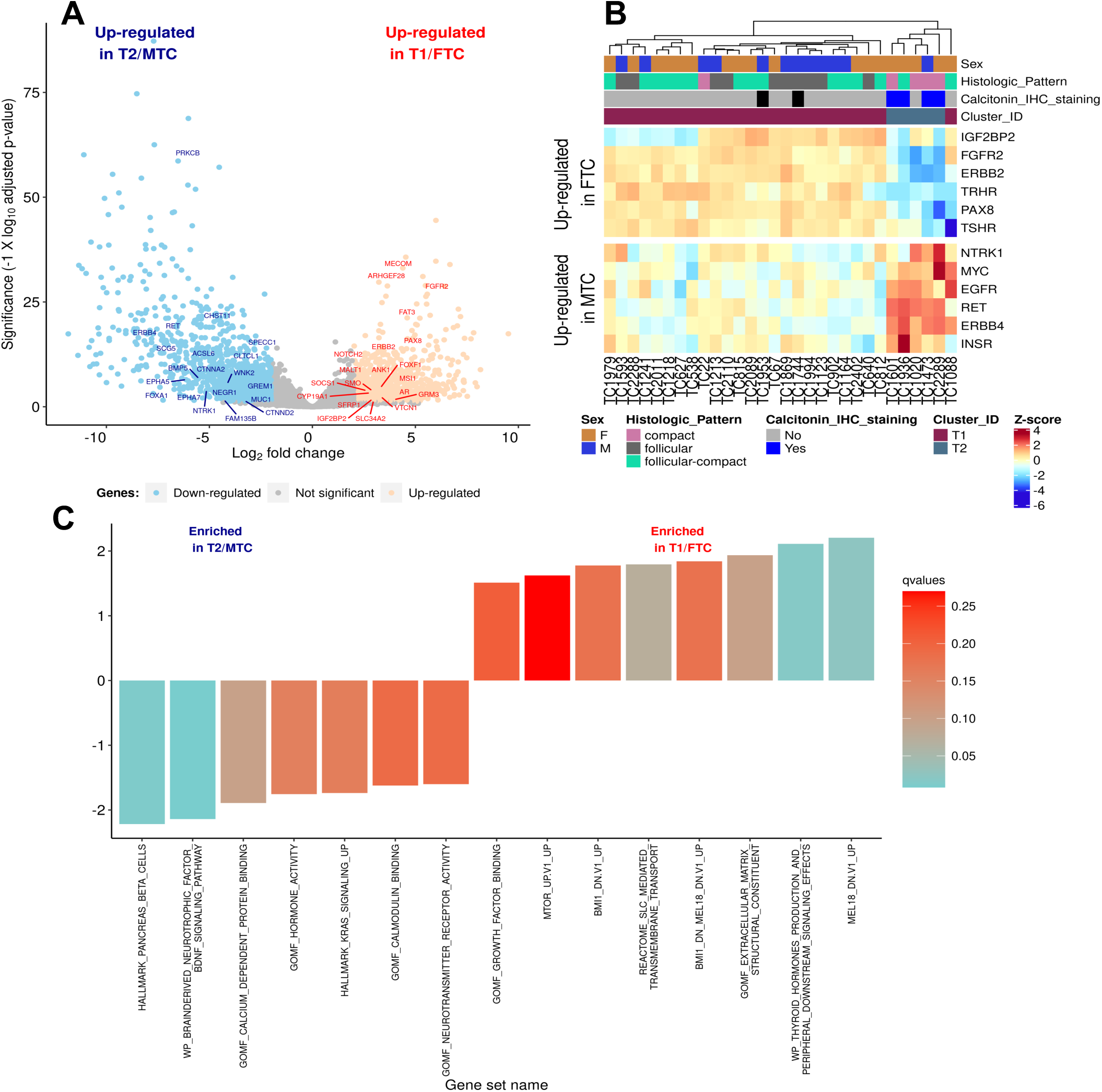
Enriched genes and pathways in canine follicular and medullary thyroid carcinomas. **A**. Volcano plot of differentially expressed genes (DEGs) between clusters. Labelled genes are differentially expressed cancer genes. The downregulated genes in the volcano plot represent genes enriched in T2 cluster (MTC). **B.** Heatmap of selected DEGs representative of each cluster. C. Significantly enriched pathways in T1 and T2 clusters. Pre-ranked differentially expressed genes were used in Gene Set Enrichment Analysis (GSEA) within the ClusterProfiler R package to identify enriched pathways. The complete list of enriched pathways is provided in S3 Table.

The enriched gene sets/pathways identified in T1/FTC group included extracellular matrix, growth factor binding, thyroid hormone production, and membrane-bound transporter gene sets (**Figure 3C**). Additionally, oncogenic signatures related to Polycomb-group genes (BMI1 DN MEL18 DN.V1 UP, MEL18 DN.V1 UP, BMI1 DN.V1 UP) were also enriched, suggesting dysregulation of epigenomic pathways in T1/FTC samples. These oncogenic signatures are also enriched in human anaplastic TC [43]. Bmi-1 supports self-renewal in stem cells by suppressing the expression of p16/INK4A and p19/ARF. The gene sets enriched in the T2/MTC group included endocrine and KRAS-related Hallmark gene sets, calcium-related gene sets, and neuron-related gene sets (**Figure 3C**).

To compare the canine and human transcriptomes we identified overlapping cross-species DEGs between tumor and normal samples. There were 3,155 and 1,230 genes differentially expressed between canine FTC and normal thyroid samples (**S5 Table**). Some of the key cancer genes up-regulated included *KIT*, *DUSP4*, *ETV4*, *INSR*, and *VEGFA*. One key tyrosine kinase gene (*ERBB2*) that was significantly up-regulated in FTC compared to MTC had a log_2_ fold change of 1.53 (adjusted p-value 0.003) in FTC compared to normal thyroid samples. Although this does not meet the log_2_ fold change >2 criteria we used for defining DEGs, this gene was significantly up- regulated in FTC samples (**S2 Fig**).

We have identified 1,932 and 939 up- and down-regulated genes, respectively, in canine MTC relative to normal canine thyroid tissues (**S6 Table**). Using the human equivalent DEG data from Minna et al. 2022, we have identified 13% gene overlap in this cross-species study [44]. Some of the key cancer genes that were up-regulated in MTC samples in both canine and human transcriptomes were *FOXA1*, *RET*, *ETV4*, *DUSP4*. The *FOXA1* gene was ubiquitously over- expressed in human MTC, but not in FTC, and might serve as a diagnostic marker [45]. Moreover, four of the top ten DEGs identified in another study of human MTCs, [14] *CALCA, CALCB, GFRA4,* and *SEMA3E,* were also significantly upregulated in canine MTCs (**S3 Fig**).

#### ERBB2 signaling in FTC samples

*ERBB2* (*HER2*) is one of the four epidermal growth factor (EGF) family members encoding a transmembrane receptor tyrosine kinase. *ERBB2* was significantly over-expressed (Kruskal-Wallis, *p* = 0.0003) in FTC tumors relative to normal thyroid and MTC tumors (**S2 Fig**). In contrast, two other EGFR family members, *ERBB4* (Kruskal-Wallis, *p* = 0.002, **S2 Fig**) and *EGFR* (Kruskal-Wallis, *p* = 0.01, **S2 Fig**), were significantly overexpressed in MTC tumors when compared to normals and FTC tumors. Additionally, HER2 IHC scores, using either the ASCO/CAP or Peña guidelines, were statistically different between FTC and MTC (Mann-Whitney, *p* = 0.0007 for both scoring schemes), with all MTC tumors considered HER2 negative (scores 0/1) while FTC tumors spanned the entire score range (0-3), the distribution of which was similar to a recent retrospective study of canine FTC [46]. The *ERBB2* expression levels observed through RNAseq were aligned with those determined by IHC scores (Kruskal-Wallis, *p* = 0.0002-0.0003, **S4 Fig**).

Given this differential expression pattern of EGF receptors in tumors and the presence of a HER2-driving mutation in the CTAC cell line [73], we assessed the enrichment of ERBB2 pathways in FTC tumors and surveyed for *HER2^V659E^* variant and HER2 protein expression. We identified several ERBB2 related pathways significantly correlated with *ERBB2* expression (**Figure 4A**). *ERBB2* expression was correlated with GSVA scores of reference ERBB signaling gene sets derived from KEGG, WikiPathway, and PID databases. Two KEGG gene sets, the ERBB2-RAS-ERK pathway and over-expression of *ERBB2* leading to PI3K signaling, were also correlated with *ERBB2* expression, suggesting possible PIK3-driven signaling in FTC samples. Similar to a previous publication that reported upregulation of PIK3/AKT pathway in canine TC [16], *EGFR* was up-regulated in MTC samples, and three of the five FTC over-expressed genes (*PIK3CA*, *AKT2*, and *PDPK1*) were up-regulated in this study as well (**S5 Fig**). Further, *ERBB2* expression was anti-correlated with the ERBB4 signaling pathway and *ERBB4* expression was anti-correlated with ERBB2 signaling pathways, emphasizing the probable effects of differential expression profile of these two genes in canine MTC and FTC tumors (**Figure 4A-B**). We also investigated the FTC samples where we correlated *ERBB2* expression to other genes, and identified the significantly associated members of the following gene sets, KEGG MEDICUS VARIANT ERBB2 OVEREXPRESSION TO PI3K SIGNALING PATHWAY and KEGG MEDICUS REFERENCE EGF ERBB2 RAS ERK SIGNALING PATHWAY. *ERBB2* was significantly correlated with PIK3-AKT signaling genes like *AKT2*, *RPS6KB2* and anti-correlated with *PIK3CA* and *PIK3CB* gene expression (**Figure 4C**). ERBB2 expression was anti-correlated with 4 of the 16 members of EGF-ERBB2-RAS-ERK signaling pathway (*SOS1*, *SOS2*, *BRAF*, and *NRAS*) and positively correlated with *GRB2* and *ARAF* expression levels (**Figure 4C**). Over-expression of ERBB2 might be attributed to genomic amplification or transcriptional regulation and is usually coupled with poor clinical outcomes in human cancers [47]. Sanger sequencing for the *HER2^V659E^* variant, identified as a homozygous variant in the CTAC canine TC cell line [48], exhibited a potential variant peak on the reverse strand chromatogram in six canine TC FFPE samples; however, no somatic variants were identified in WES or RNAseq of matching frozen samples. Based on the depth of reads of the genomic sequencing analyses, we feel that this mutation is not present in this cohort. The CTAC cell line was established in 1964 [49], and since that time, it is possible that there has been preferential selection of subclones that harbor this mutation, rendering it homozygous.

**Fig 4.**
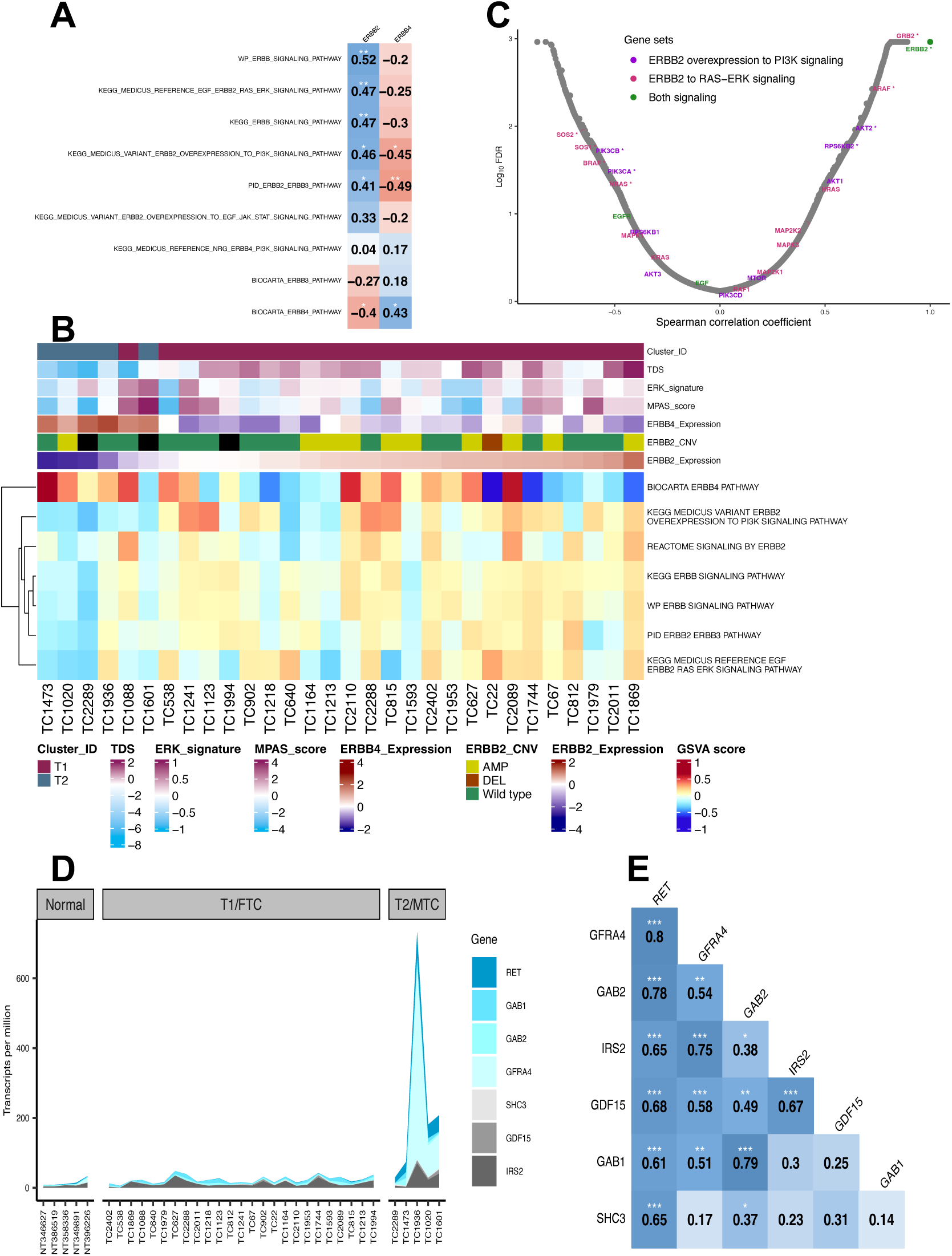
EGF receptor family and *RET* signaling genes in canine thyroid carcinomas. **A-C**. ERBB2 signaling in canine thyroid carcinomas. **A**. Spearman correlation coefficients (*p-val <0.05, **p-val <0.01), of GSVA scores derived from ERBB-related gene sets and *ERBB2*/*ERBB4* expression values. **B**. Heatmap of selected pathway GSVA scores and its association with *ERBB2*, *ERBB4* expression levels, *ERBB2* copy number variants (CNV) derived from Sequenza (Seqz) bioinformatics resource, thyroid differentiation score (TDS), ERK signature, and MPAS score. The samples were arranged in ascending order of ERBB2 expression levels. **C**. Scatterplot of Spearman coefficients and Log_10_ FDR derived from the correlation between *ERBB2* and 15,015 gene expression levels in FTC samples. The members of two signaling gene sets are highlighted in two different colors. The members common between the two gene sets are highlighted in green. **D-E**. RET signaling in thyroid carcinomas. **D**. Area plot displaying over-expression of seven RET signaling genes in MTC tumors. **E**. The Pearson correlation matrix of genes shows that *RET* expression was significantly correlated with genes that encode their ligands, adaptors, co-receptors, and activators.

In all, the protein and gene expression of HER2/*ERBB2* in this cohort poses an interesting molecular marker and potential therapeutic target for canine TC, as discussed later.

#### RET signaling in MTC samples

Receptor tyrosine kinase, *RET* or *MEN2A/B*, gene expression was significantly up-regulated in MTC tumors, suggesting that RET could drive these tumors in dogs. However, unlike humans, no *RET* driver mutations or fusions were identified in canine MTC. Based on RET signaling in neurons [50], we have identified a set of ligands, RET co-receptors, and downstream pathway activators that significantly correlated with *RET* transcript expression (**Figure 4D-E**). Of the five known RET ligands [51], only GDF15 was significantly up-regulated in T2/MTC samples. GDF15 binds to GFRAL, which is a co-receptor of RET and can activate phosphorylation of ERK in the presence of wild-type RET [52]. In addition to RET and GDF15, multiple genes that are components of the RET signaling pathway were significantly correlated with RET expression and up-regulated in MTC samples. One of these genes, co-receptor GFRA4, directly binds with RET to form a functional receptor for PSPN (persephin), thereby activating intracellular RET signaling [53]. Further, RET signaling adaptors, *GAB1*, *GAB2* and *IRS2*, were also up-regulated in canine MTC, which in turn recruits SH2 domain-containing proteins to activate kinase signaling pathways [54]. In this study, the expression of neuron-specific gene *SHC3* (*Rai*) was correlated with *RET* expression and was significantly up-regulated in T2/MTC group compared to both the normal and T1/FTC groups (**Figure 4D**). Although it is difficult to assess the specific downstream pathway of RET signaling in canine MTC from transcriptomic data alone, the relatively high expression of RET and its signaling components suggests activity of this pathway in medullary tumors. In humans, overexpression of wild-type *RET* has been observed in several cancer types, including breast cancer and pancreatic ductal carcinomas, and is associated with high-grade metastatic tumors [55]. Germline mutations in human *RET* are responsible for the autosomal dominant genetic syndromes multiple endocrine neoplasia type 2 (MEN2) and familial MTC (FMTC). These syndromes are characterized by an increased incidence of MTC, as well as an increasing risk for the development of pheochromocytoma, primary hyperparathyroidism, and other endocrine neoplasms. While only approximately 25% of human MTCs are hereditary, as many as 75% of sporadic human MTC have somatic mutations in the RET oncogene [56].

#### Mutational spectrum

Whole exome sequencing of 27 tumors and matched normals [mean depth: 220X (±27.8X) for tumors and 100.8X (±10.9X) for normals] identified a somatic tumor mutational burden (TMB) of 2.9±2.2 non-synonymous protein coding mutations per Mb sequenced, similar to the human PTC TMB (**Figure 5A**) [11]. However, compared to other canine cancers like osteosarcoma, hemangiosarcoma and melanoma, where median TMB was ≥ 1 mutations per Mb, the TC TMB was relatively higher [57]. Unlike human cases, the mutational burden did not correlate with age (Pearson correlation, *p* = 0.8), weight (Pearson correlation, *p* = 0.9), tumor differentiation status (Kruskal-Wallis, *p* = 0.22), TDS (Pearson correlation, *p* = 0.09), or progression-free interval (PFI) (COXPH, *p* = 0.4). However, dogs with low TMB (i.e., low somatic mutations) were associated with shorter PFI (< 250 days) (Kruskal-Wallis, *p* = 0.025, **S6 Fig**).

**Fig 5.**
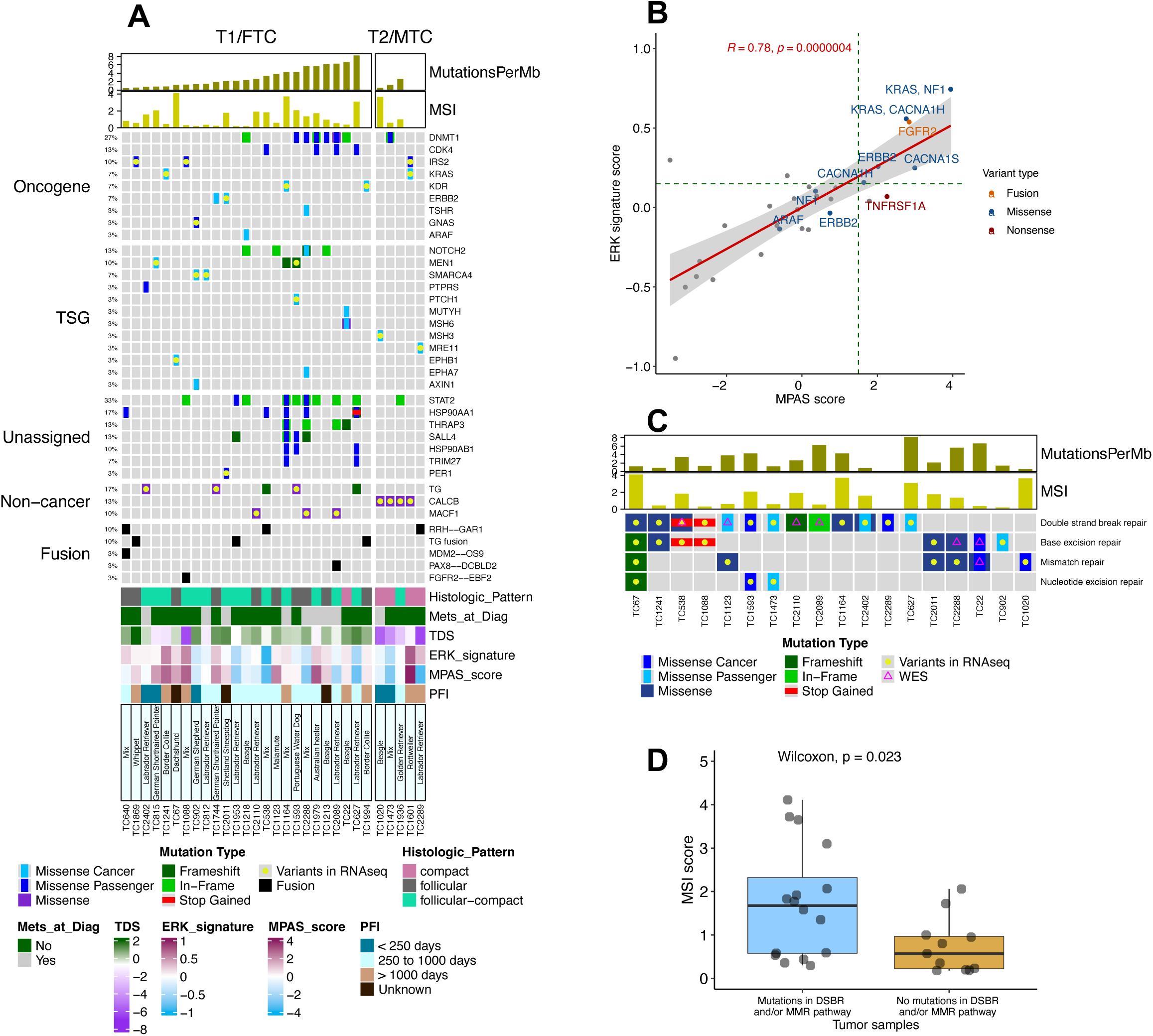
Mutational landscape of canine thyroid tumors. A. An Oncoplot of variants identified from WES and/or RNAseq variant calling pipelines. The cancer genes mutated in at least 10% of samples, predicted cancer-driver genes, and thyroid-related gene variants were plotted here. B. Scatterplot showing the MPAS and ERK score distribution across 30 tumor samples, showing a strong correlation between two pathway scores across 30 samples. Additionally, samples with mutations in genes related to these two pathways were associated with high MPAS and ERK scores. C. Illustration of samples with mutations in four different DNA repair pathways. These variants were identified either from WES and/or RNAseq pipelines. There were no signification correlations between Mutations per MB and mutations in DNA repair pathway genes. These genes are listed in Table 3. D. Boxplot illustrating the distribution of MSI scores, grouped by samples with and without DSBR and MMR mutations. Samples with double-strand break repair (DSBR) and/or mismatch repair (MMR) gene variants exhibited a significantly higher MSI score than non-mutant cases.

Relative frequencies of DNA substitutions classified by the conventional 96 mutation types were used to construct three *de novo* mutational signatures from 27 TC [58]. The most common type of substitution was C < T in both FTC and MTC cohorts (**S7A Fig**). Of the three *de novo* mutational signatures, two were most similar to SBS5 and SBS26, respectively, and one was a novel signature (SBSA) (**S7B Fig**). The etiology of SBS5 is unknown but correlates with age, and the SBS26 signature is associated with the DNA mismatch repair pathway. Our data show a statistically significant association between SBS26 and mutational burden (Kruskal-Wallis, *p* = 0.0009, **S7C Fig**). Pan-cancer analysis of human tumors has shown that the mutational burden in TC is low and dominated by clock-like SBS signatures associated with aging (SBS1 and SBS5) [58]. Less frequently, signatures SBS2 and SBS13 associated with APOBEC activity have been identified. Signature SBS13 serves as a predictor of resistance to radioiodine therapy [59].

#### Mutational landscape distinctly different from human counterpart

We have identified cancer variants in both MTC and FTC tumors using WES and RNAseq data mapped against the CanFam3.1 genome. Following variant filtering, 2,587 protein-coding variants within 1,581 genes were identified from WES of 27 samples and 1,930 variants in 1,429 genes from RNAseq of 30 samples (**S7 and S8 Tables**). The top cancer genes with recurrent variants included seven genes: *STAT2* (33%), *DMNT1* (27%), *HSP90AA1* (17%), *CDK4*, *NOTCH2*, *THRAP3*, and *SALL4* (13% each) (**Figure 5A**). The variants in these genes included 24 SNV, of which only one was predicted by FATHMM to be a cancer driver (*NOTCH*^C1490W^), and 16 INDELs. However, the mean allelic frequency of these variants (n=66) was very low, 3.3% ± 3.2%, with a median sequencing depth of 228X. As such, the functional consequences of these variants on cancer progression were difficult to predict. This profile distinctly differs from human TC, where recurrent driver mutations were observed in *BRAF*, and RAS-family genes [11]. Of the MAPK pathway genes that are frequently mutated in human TC, we have identified one gene with driver mutation, *KRAS^Q155R^*, in two samples, including one FTC (TC1241) and one MTC (TC1601). Additional MAPK pathway gene variants like *ERBB2^H42R^ ^&^ ^T323M^*(TC1744 and TC2011), *ARAF^T235P^* (TC1218), *NF1^D1430N^ ^&^ ^L928P^* (TC22, TC1601), *TNFRSF1A^W406^\** (TC1123), two calcium channel subunit genes (TC67, TC1979), and an *FGFR2* fusion (TC1088) were also identified. About 67% of these variants (8 out of 12) were also associated with relatively high MPAS and ERK signature scores (>75^th^ percentile), suggesting a possible increase in MAPK pathway activity relative to other tumors within this cohort (**Figure 5B**). In addition, there was a high concordance between the MAPK score and ERK score in these samples (Pearson correlation coefficient = 0.78, *p* < 0.001), which is plausible since ERK signaling is part of the broader MAPK signaling pathway.

The next TC-relevant gene identified was a tumor suppressor gene, Multiple endocrine neoplasia type 1 or *MEN1*, with three variants (A100P, N282Tfs*86, Y312Lfs*5) in FTC (**Figure 5A**). In humans, germline *MEN1* mutations have been associated with a hereditary cancer syndrome characterized by pituitary, parathyroid, and pancreatic adenomas [60], and somatic variants were linked to <1% of TC (https://bit.ly/3ZiGDaR). In our dataset, the 10% of tumors with somatic *MEN1* mutations were accompanied by concomitant decreased expression of *MEN1* transcripts compared to samples with wild-type *MEN1* (**S8 Fig**). As a tumor suppressor gene, the frameshift mutations in *MEN1* were probable drivers and the missense mutation was a predicted cancer driver in these canine patients.

The German shepherd dog patient (TC902) carried a *GNAS*^A204D^ variant, previously identified as a likely novel recurrent driver in Dutch German longhaired pointer dogs [18]. However, this variant was predicted by FATHMM to be a cancer passenger mutation (A249D), suggesting in humans this alteration might not be a cancer driver. Hotspot activating mutations (R201 and Q227) and amplifications of GNAS have been identified in human tumor types including pituitary, pancreatic, colorectal, and TC [61]. This heterogeneity in the tumor mutational landscape is distinctly different from the human TC counterpart. However, similar genetic heterogeneity has been seen in human neuroendocrine tumors [62].

We also observed histotype-related mutations in two genes: *TG* in 5 of 25 FTC and *CALCB* in 4 of 5 MTC samples. Three additional canine samples carried *TG* fusions, which led to an aggregate of 32% of patients with *TG* variants. Although TG levels were not significantly increased in the samples with mutations compared to wild-type *TG* samples, samples with *TG* fusions had relatively higher levels of *TG* expression (**S9A Fig**). Somatic variants in TG have been reported in about 4% of human PTCs which might result in dysregulation of thyroid hormone synthesis [11]. In addition to *TG*, another FTC histotype-related variant identified was a putative driver mutation in *TSHR* (TC2288). In the case of the MTC samples, *CALCB*^A93G^ variants in 4 tumors were associated with higher expression levels of the calcitonin B gene compared to the one sample (TC2289) that lacked that mutation (**S9B Fig**).

The majority of the variants identified from mapping against Canfam3.1 were also identified from Canfam4 (See **S1 Appendix**). One of the top recurrent cancer genes identified from mapping to CanFam4 was *MUC4*; however, only some of these variants could be validated via Sanger sequencing due to the highly repetitive sequence of the *MUC4* gene. Taken together, we see a diverse and heterogeneous landscape of mutation in canine FTC and MTC, which contrasts with their human counterparts.

#### Fusion genes in canine thyroid carcinomas

In addition to short variants, we plotted selected gene fusions that were identified from mapping RNASeq data against both CanFam3.1 and CanFam4 genomes (**S9 Table**). These results also encompass a heterogeneous population of fused genes that were mutated in less than 10% of the canine tumors. Putative driver fusions in canine FTC samples FGFR2--EBF2 and PAX8-- DCBLD2, include gene partners *FGFR2* (VCL-FGFR2) and *PAX8* (PAX8-PPARG) that have been identified in human TC [20]. These fusions led to elevated expression of the downstream partner, but the functional implications of this are unclear as these partners are not considered cancer genes. The most recurrent fusion identified was RRH-GAR1 in 3 tumors. Although its functional significance is unknown, the fusion was associated with elevated expression in 2 of the 3 samples (**S10 Fig**). The *RRH* or *Peropsin* gene encodes a G-protein coupled receptor and *GAR1* gene encodes small nucleolar ribonucleoprotein. However, the most common recurrent fusion gene in human TC, *RET*, was not identified in this cohort of canine TC [63]. As mentioned earlier, we have identified three samples with TG fusions in FTC samples, which were validated using Sanger sequencing.

In addition to the fusions identified using both CanFam3.1 and CanFam4, potentially relevant fusions found only with the CanFam3.1 mapping and associated with elevated transcripts were MDM2-OS9 (TC640) and GNAS fusions (TC1593, TC2289, TC2402). GNAS transcripts were significantly elevated in TC2289 and TC1241 compared to the rest of the cohort.

#### Impaired DNA repair pathways in canine TC

Leveraging variant calling from both WES and RNASeq data, we identified mutated DNA repair pathway (DRP) genes in 60% of canine TC (**Figure 5C**). This included 24 genes with 26 variants that are functional components of mismatch repair (MMR), base excision repair (BER), nucleotide repair (NER), and double strand break repair (DSBR) pathways (**Table 3**). About 40% of these variants from the RNAseq variant calling pipeline were also identified in the matched normal WES sample, suggesting a probable germline variant. Although we have taken measures to remove the majority of germline variants from RNAseq data using the Panel of Normals and public resources, sample-specific variants were not filtered from RNAseq data. We used two different genomic scores to find an association with these DRP mutations: MSI score (the percentage of unstable microsatellite loci within the tumor genome), and mutational burden (the fraction of somatic mutations per megabase sequenced). The canine patients with DSBR and/or MMR pathway mutations (n=17) had significantly higher MSI scores compared to the rest of the cohort (n=13) (Wilcoxon, *p* = 0.023, **Figure 5D**). However, there was no significant association between DRP mutations and mutational burden. In addition, variants in MMR pathway genes, *MSH3*, *MSH6*, *LIG1*, *MCM9*, and *POLD3*, were associated with either relatively high MSI score and/or high mutational burden in 6 tumor samples. The sample with the highest mutational burden (TC627) carried an *ATM* cancer variant, a component of DSBR pathway. The inactivation of genes in DNA repair pathways may contribute to the heterogeneous mutational landscape observed in canine TC, and in turn facilitate genomic instability and tumor progression.

**Table 3.** DNA repair pathway genes mutated in canine thyroid tumors.

| Gene Name | Sample ID | HGVSp | Sequencing Pipeline | Repair pathway | IGV validation |
| --- | --- | --- | --- | --- | --- |
| DCLRE1C | TC2110 | G207Efs*20 | WES | DSBR | RNASeq; Tumor WES |
| POLD3 | TC67 | A362Sfs*5 | RNA | BER, DSBR, MMR, NER | RNASeq |
| BARD1 | TC2089 | Q542del | WES | DSBR | Tumor WES |
| ACD | TC2288 | V266I | WES | BER | RNASeq |
| POT1 | TC902 | M711L | RNA | BER | RNASeq |
| SIRT6 | TC1241 | R145H | RNA | BER, DSBR | RNASeq; Tumor and Normal WES |
| LIG1 | TC2011 | S444R | RNA | BER, MMR | RNASeq |
| MUTYH | TC22 | R658W | WES | BER, MMR | RNASeq; Tumor WES |
| ATM | TC627 | K687Q | RNA | DSBR | RNASeq |
| BARD1 | TC1123 | L654F | WES | DSBR | Tumor WES |
| KAT5 | TC67 | I531L | RNA | DSBR | RNASeq; Tumor and Normal WES |
| MDC1 | TC1164 | R1649Q | RNA | DSBR | RNASeq; Tumor and Normal WES |
| MRE11 | TC2289 | L286V | RNA | DSBR | RNASeq |
| MUS81 | TC538 | H375N | WES | DSBR | Tumor WES |
| NSD2 | TC2402 | A1309T | RNA | DSBR | RNASeq; Tumor and Normal WES |
| RIF1 | TC2402 | I2399L | RNA | DSBR | RNASeq |
| ERCC4 | TC1473 | H201Q | RNA | DSBR, NER | RNASeq; Tumor and Normal WES |
| TP53 | TC1593 | I247L | RNA | DSBR, NER | RNASeq |
| MCM9 | TC2288 | R450K | RNA | MMR | RNASeq; Tumor and Normal WES |
| MLH3 | TC1123 | S1103Y | RNA | MMR | RNASeq; Tumor and Normal WES |
| MSH3 | TC1020 | M282R | RNA | MMR | RNASeq; Tumor and Normal WES |
| MSH6 | TC22 | E998D | WES | MMR | Tumor WES |
| MSH6 | TC22 | R990H | WES | MMR | Tumor WES |
| NEIL1 | TC1088 | W296* | RNA | BER | RNASeq; Tumor and Normal WES |
| POLQ | TC538 | R793* | RNA | BER, DSBR | RNASeq; Tumor and Normal WES |
| TIMELESS | TC1088 | Q1283* | RNA | DSBR | RNASeq; Tumor and Normal WES |

#### Survival analyses

Many dogs in this cohort experienced prolonged survival post-thyroidectomy with a median PFI of 1837 days (5.03 years) and median ST of 1892 days (5.18 years) for all TC. Among the censored dogs (i.e., those which did not have detectable disease, nor death due to disease) with TC, the median PFI (excluding dogs with metastasis at diagnosis) was 761 days (2.08 years; range 0–3138 days) and the median ST was 778 days (2.13 years; range 28–3138 days). Similarly, the median PFI was 1837 days and the median ST was 1892 days for dogs with FTC, while the median PFI and ST were not reached for dogs with MTC due to patient censoring (**Table 1, S11 Fig**). The outcomes were not significantly different among dogs with FTC vs MTC, which has been previously reported [9]; however, the MTC cohort in this study was underpowered (n=6) and future studies are needed to clarify the clinical distinction of FTC vs MTC. Currently, canine TC are treated similarly, and the distinction of FTC vs MTC is not routinely sought, presumably based on the previous study where similar outcomes were reported regardless of origin of TC. Thyroidectomy is the treatment of choice for amenable tumors with use of adjunctive therapy post- operatively not demonstrating consistent clinical benefit in the canine literature [29, 64]. In this cohort, 30% of dogs (n=18) underwent adjunctive therapies (i.e., chemotherapy, radiation, and/or immunotherapy) post-thyroidectomy. Six of these dogs had metastasis at diagnosis (n=13 for all dogs presenting with metastasis at diagnosis), and seven developed metastases after diagnosis. Full chemotherapy protocols were unavailable for review but included single agent carboplatin (n=10), single agent doxorubicin (n=3), combination carboplatin/doxorubicin (n=2), combination melphalan/doxorubicin (n=1), and combination doxorubicin/carboplatin/vinorelbine (n=1). Two dogs received palliative radiation therapy in addition to chemotherapy (single agent carboplatin and multiagent carboplatin/doxorubicin/vinorelbine), and one dog received toceranib (Palladia®) a tyrosine kinase inhibitor, as a single agent. The dogs that received post-thyroidectomy adjunctive therapy experienced shorter PFI and overall survival compared to dogs that had thyroidectomy (surgery) alone (median PFI 961 days vs median not reached due to patient censoring, *p* = 0.0024, HR 5.1, 95%CI 1.4-18.0; median ST 1892 days vs not reached due to censoring, *p* = 0.0214, HR 3.8, 95%CI 0.96-15.17; **Figure 6A-B**). A similar trend was seen when thyroidectomy alone was compared to those receiving only chemotherapy as adjunctive (median PFI 1057 days vs not reached, *p* = 0.012, HR 4.28, 95%CI 1.04-17.59, median ST not reached vs 1057 days, *p* = 0.012, HR 0.26, 95%CI 0.07-0.94; **S11 Fig**); however, overall survival was not statistically different between dogs receiving surgery only and surgery plus chemotherapy (1892 days vs not reached due to patient censoring *p* = 0.099, HR 2.94, 95%CI 0.60-14.47). Since nearly 50% of dogs presenting with metastasis at diagnosis underwent adjunctive treatment, these therapies were likely recommended due to perceived negative prognostic findings so direct causation of these therapies to outcome is not clear [65]. Metastasis at diagnosis was not considered a negative prognostic factor in our cohort, which is supported and conflicting with other studies (**S11 Fig**) [29, 65]; as such, it remains unclear the significance of this finding in canine TC.

**Fig 6.**
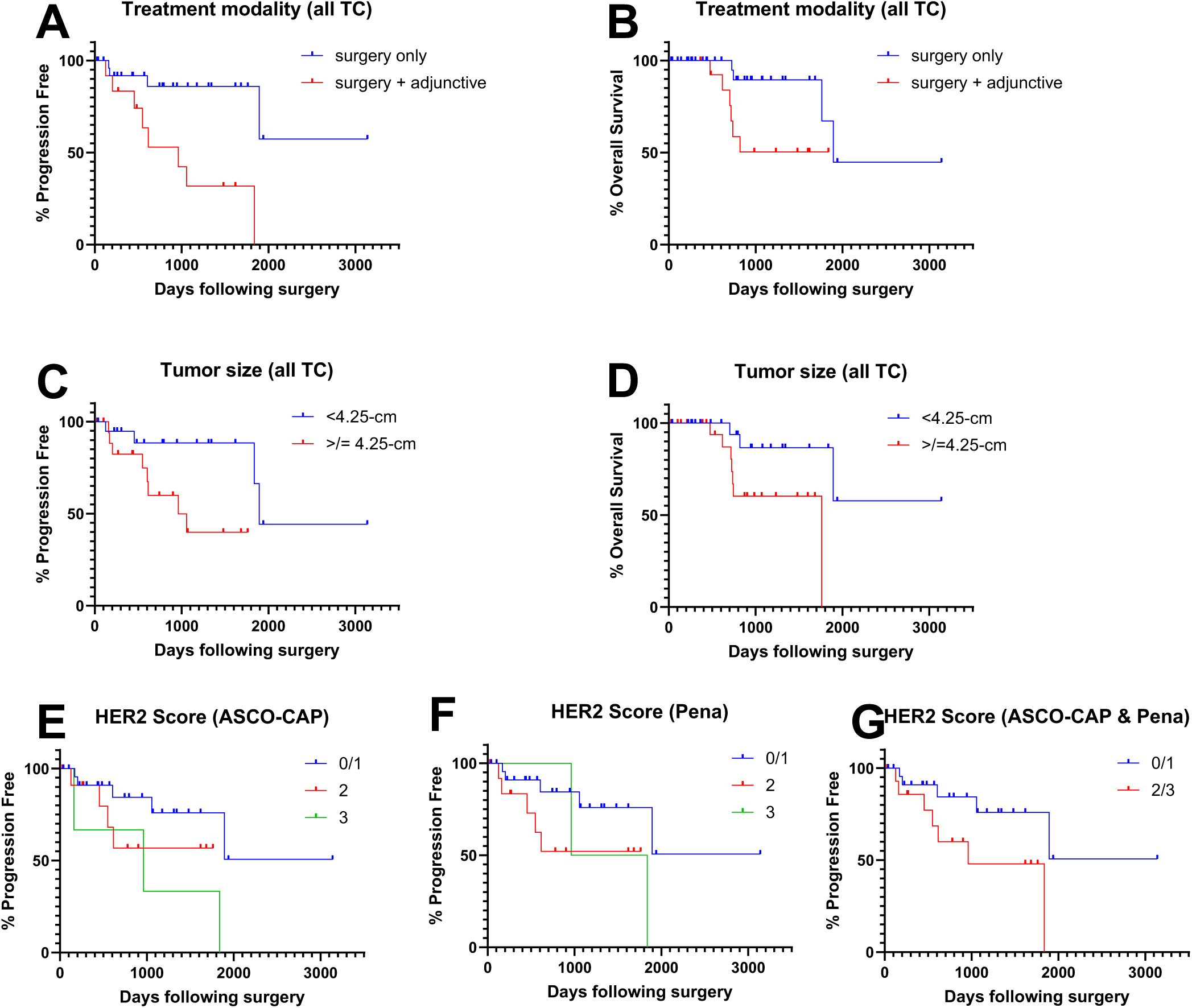
Survival analyses. **(A-B: Treatment modality)** Median PFI of dogs that received surgery only was not reached due to patient censoring, while for dogs receiving surgery and any adjunctive treatment (chemotherapy, radiation, and/or immunotherapy) was 961 days (2.63 years, *p* = 0.0024, HR 0.19, CI95% 0.05-0.69) What about overall survival?. **(C-D: Tumor size)** Median PFI of dogs with thyroid tumors less than cohort median size (<4.25-cm) was 1892 days (5.18 years) vs 961 days (2.63 years) for dogs with tumors ≥4.25-cm; these are considered statistically different (*p* = 0.0216, HR 0.29, CI95% 0.09-0.94). Median ST of dogs with thyroid tumors <4.25-cm was not reached due to patient censoring vs 1761 days (4.82 years) for dogs with tumors ≥4.25-cm; these are considered statistically different (p = 0.025, HR 0.25, CI95% 0.07-0.92). **(E-G: HER2 IHC score)** Median PFI of dogs with ASCO/CAP HER2 Scores of 0/1 (negative) was not reached due to patient censoring, while dogs with Score 3 was 961 days (2.63 years); these were considered statistically different (*p* = 0.019, HR 0.22, CI95% 0.02-1.90); this finding was not observed with the Pena scheme. However, when combining HER2 Scores of 2 and 3 (which was identical for ASCO/CAP and Pena schemes), median PFI was 1837 days (5.03 years) and was considered statistically different than dogs with Scores 0/1 (p = 0.046, HR 0.34, CI95% 0.10-1.13).

Increased tumor diameter was associated with decreased ST and PFI based on a median split of all TC (4.25 cm; **Figure 6C-D**). Animals with smaller tumors experienced a median PFI of 1892 days (5.18 years) compared to 961 days (2.63 years) for animals with larger tumors (*p* = 0.0216, HR 0.29, 95%CI 0.09-0.94). The median ST of animals with smaller tumors was not reached due to patient censoring, while the median ST of animals with larger tumors was 1761 days (4.82 years) (*p* = 0.025, HR 0.25, 95%CI 0.07-0.92). Similar findings were detected among dogs with FTC, in which dogs with tumors <4.25 cm (median split of FTC) experienced a statistically longer PFI (median 1892 days vs 961 days; *p* = 0.0217, HR 0.29, 95%CI 0.09-0.94) and ST (median not reached vs 1761 days; *p* = 0.024, HR 0.25, 95%CI 0.07-0.91) than dogs with tumors ≥4.25 cm. Correlations between larger tumor size and poor outcome have been previously reported for canine TC but are not consistently linked in the literature [9, 23, 29, 34, 65, 66]. Smaller tumors may be more amenable to complete excision and earlier excision could prevent or delay metastasis, as well as other complications from tumor growth and invasion.

HER2 IHC score was inversely correlated with PFI for all TC (**Figure 6E-G**). In particular, dogs with tumor ASCO-CAP score 3 had statistically different PFI than those with 0/1 (961 days vs not reached due to patient censoring, *p* = 0.019, HR 4.6, 95%CI 0.53-40.27). This difference was not appreciated when the Pena scheme was applied for score 3 vs 0/1 (1399 days vs not reached due to patient censoring, *p* = 0.145, HR 3.09, 95%CI 0.30-31.57). However, when scores 2 and 3 were combined (which made data sets for the ASCO-CAP and Pena schemes identical), the median PFI was 961 days versus not reached for scores 0/1, which were statistically different (*p* = 0.046, HR 2.96, 95%CI 0.88-9.96). HER2 IHC scores did not have any correlations with overall survival. As previously mentioned, HER2 is a member of the receptor tyrosine kinase family and signals downstream through mitogen-activated protein kinase (MAPK)/ERK and PI3K/Akt pathways, which promote cellular proliferation, survival, invasion, and migration. Over-expression of *ERBB2* might be attributed to genomic amplification or transcriptional regulation and is usually coupled with poor clinical outcomes in human cancers [47]. In canine mammary carcinomas, HER2 overexpression (IHC scores 3 +/- 2) has been correlated with longer PFI and overall survival but also with presence of higher histologic grade and even proliferative index, although these are inconsistently associated in the literature [67–72]. As such, the diagnostic and prognostic utility of HER2 IHC remains to be clarified in the veterinary literature and future directed studies are needed.

The remaining clinicopathologic parameters (i.e., age at diagnosis, weight at diagnosis, tumor location, thyroid hormone status, tumor histotype, differentiation, nuclear atypia, and presence of microscopic tumor emboli, necrosis, hemorrhage, or bone/mineral) were not associated with outcome.

## Materials and Methods

### Sample acquisition and processing

The canine thyroid tumors were acquired from the Colorado State University (CSU) Flint Animal Cancer Center Biorepository. All tumors had formalin-fixed, paraffin embedded (FFPE) tissue and a subset also had frozen tissue sections. FFPE tumor tissue blocks were routinely processed for hematoxylin and eosin (H&E) staining and evaluated by two veterinary pathologists (SNS & DPR) to confirm adequate tumor sample for inclusion in pathologic and genomic analyses. Both DNA and RNA were extracted from each available frozen tumor sample using TRIzol (Invitrogen, Catalogue # 15596026) according to the manufacturer’s protocol. Matched normal sample DNA was also extracted for whole exome sequencing (WES). The DNA and RNA samples were purified using DNeasy or QiaAMP DNA Blood mini kits and RNeasy (Qiagen, Catalogue #69504, 51104, 74004), respectively. RNA/DNA purity, quantity and integrity were determined by NanoDrop (ThermoFisher Scientific) and TapeStation 4200 (Agilent, CA, USA) analysis prior to DNA and RNA-seq library preparation. The Zymo-Seq RiboFree Total RNA Library Kit was used (Zymo Research, CA, USA, Catalogue # R3000) with an input of 300 ng of total RNA to generate RNAseq libraries. Paired-end sequencing reads of 150 bp were generated on a NovaSeq 6000 (Illumina Inc., CA, USA) sequencer at a target depth of 60 million paired-end reads per sample. For WES, the SureSelectXT Target Enrichment System for Illumina Paired-End Multiplexed Sequencing Library kit was used to create genomic DNA libraries that were also sequenced on a NovaSeq6000 generating 150 bp paired-end reads. Raw sequencing reads were de-multiplexed using bcl2fastq. Sequencing and library prep for DNA and RNA was carried out at the Genomics Shared Resource, University of Colorado Anschutz Medical Campus (RRID: SCR_021984). In addition, we downloaded RNAseq fastq files of normal thyroid tissues from an open-access dog epigenomic resource, BarkBase (https://barkbase.org/, BioProject PRJNA396033) [73].

#### Clinicopathological data

Medical records were reviewed to identify clinical data including age at diagnosis (i.e., surgery), sex (including neuter status), weight, breed, tumor localization, largest tumor diameter (cm), and thyroid hormone testing (if available). Survival data (progression free interval, PFI, and overall survival, ST; determined as time from surgery to disease progression, death due to disease, or last known follow-up) were determined from CSU medical records and/or contact with referring veterinarians.

The FFPE tumor samples were subjected to histopathologic characterization (performed on H&E-stained slides) and immunohistochemical (IHC) staining for calcitonin and HER2 by two board-certified veterinary pathologists, with consensus for each evaluated parameter. Histopathologic factors assessed were: histologic pattern (i.e., follicular, follicular-compact, and compact, based on the WHO scheme [74]); degree of differentiation (i.e., well, moderate, and poor); degree of nuclear atypia (i.e., mild, moderate, or marked); and presence of microscopic invasion/infiltration, tumor emboli, necrosis, hemorrhage, and mineral/bone. For IHC, slides were stained on the BOND Rxm system using the BOND Polymer Refine Detection kit (Leica Biosystems Inc.) with the following steps: 1) antigen retrieval with Tris-EDTA buffer, pH 9.0, for 20 minutes at 100°C (BOND Epitope Retrieval 2, Leica), 2) incubation at ambient temperature for 15 minutes with primary rabbit polyclonal antibodies for calcitonin (Abcam, ab8553; 1:200 dilution, validated based on immunoreactivity in canine thyroid (normal and carcinoma) tissue specimens – personal communication with Kansas State Veterinary Diagnostic Laboratory) or HER2 (Dako, A0485; 1:400 dilution, previously validated in canine cells/tissue [75]) diluted in BOND Primary Antibody Diluent (TBS), and 3) incubation with BOND Polymer at ambient temperature for 25 minutes, exposed to chromogen (DAB) and counterstained with hematoxylin. Tissue samples were considered calcitonin positive if the majority of tumor cells showed strong cytoplasmic reactivity. The HER2 samples were graded based on American Society of Clinical Oncology–College of American Pathologists (ASCO-CAP) [76, 77] and Peña [78] recommendations for human and canine specimens, respectively, where grade 0-1 are negative, 2 is equivocal, and 3 is positive (**S1 Fig**).

#### Processing and analysis of sequencing data

The paired end RNAseq reads (range: 51.9 – 116.3 million) were mapped against CanFam3.1 (Ensembl version 99) and CanFam4 (UU_Cfam_GSD_1.0, NCBI RefSeq assembly: GCF_011100685.1), with STAR using -genecount parameters (v 2.6.1a) [79]. The new canine genome (CanFam4) was also supplemented by three Y chromosome sequences from a Labrador retriever (ROS_Cfam_1.0, GCF_014441545.1) assembly [80]. Gene count data was normalized with DESeq2 (v1.26.0) median of ratios method [81]. Additionally, transcripts per million (TPM), a method for normalizing gene expression levels, was calculated for each gene using TPMCalculator [82]. Normalized count data was log-transformed and scaled for plotting and downstream analyses. The WES paired reads were also mapped against both CanFam3.1 and CanFam4 genomes using BWA. Prior to variant calling, the BAMs were processed in accordance with GATK best practices [83].

#### Clustering approach for sample categorization

The Uniform Manifold Approximation and Projection (UMAP) dimension reduction algorithm was used to group tumor and normal samples into distinct clusters [38] using DEseq2 log- transformed normalized data as input. A subset of genes with mean expression log_2_ > 2 and mean variance log_2_ > 6 (n = 1,013) were used for clustering samples to reduce noise and focus on biologically relevant features.

#### Differential expression and pathway enrichment analysis

DESeq2 was used to run differential expression (DE) analyses between the two canine thyroid cancer tumor clusters [81]. The sample TC1088 was identified as an outlier and was eliminated from DE analysis. For the DEG analysis pipeline, raw gene counts were filtered to retain only genes with at least 10 reads in at least 5 samples. Following the median of ratios count normalization, DESeq was used to identify differentially expressed genes (DEGs) with log_2_ fold change >2 and adjusted p-value <0.05. The DEGs were also identified between normal thyroid samples and each tumor cluster. Enriched pathways (FDR <0.25) in each cluster were identified using pre-ranked Gene Set Enrichment Analysis (GSEA) through the ClusterProfiler package in R, which employs the Kolmogorov-Smirnov goodness-of-fit test [84].

#### Gene fusion identification and processing

STAR-fusion was used to identify fusion genes from CanFam3.1 and CanFam4 genome assemblies [85]. Two canine gene fusion databases were created from both genome assemblies using prep_genome_lib.pl script from Star-Fusion. Using fastq reads as input, gene fusions were identified by the STAR-fusion tool. The fusions were filtered and manually curated to identify putative drivers. We used the following criteria to eliminate potential false-positives: a) fusion genes where both partners are from the same gene family (e.g.: PBX2-PBX3), b) fusions without official gene names, c) fusions identified in >80% of the samples, and d) fusions that were identified in both TC and normal thyroid tissues. The fusion genes were further curated to identify fusions with known cancer genes (OncoKB database). Selected fusion genes were also validated via Sanger sequencing. The forward and reverse primer sequences were reported in S2 Appendix.

#### Signaling and thyroid differentiation score

We used gene expression data to calculate the MAPK Pathway Activity Score (MPAS), ERK Score, and Thyroid Differentiation Score (TDS) [11, 86, 87]. For details see **S2 Appendix.**

#### Somatic variant calling

Duplicate reads were marked with Picard tools (v1.119) and somatic variants were called using GATK Mutect2 (v4.1.2.0) [83]. Variants were called from both RNAseq and WES data. For the WES variant calling pipeline, germline variants were eliminated using three different resources: a) matched normals, b) a panel of normals created from 118 in-house normal tissue or PBMC samples using the GATK pipeline, and c) an external germline resource identified from 722 dogs (∼90 million population variants) [88]. The remaining variants were processed using the filterMutectCalls GATK function, and variants with a PASS notation in the FILTER column were characterized as somatic variants. For the RNAseq variant calling pipeline, variants were identified using GATK HaplotypeCaller. Germline variants were filtered out following the same approach as the WES pipeline, with the exception that a matched normal was not used. Additional filtering criteria included a read depth (DP) greater than 10, an alternate allele count greater than 5, removal of splice variants, and selection of variants with an allelic frequency between 0.25 and 0.85. The VCF files with somatic variants were converted to MAF (Mutation Annotation File) format (https://docs.gdc.cancer.gov/Data/File_Formats/MAF_Format/) using the perl code: vcf2maf.pl (https://github.com/mskcc/vcf2maf). Each variant was mapped to only one of all possible gene transcripts using the “canonical” isoform from Ensembl database (v99). The variants within cancer genes were curated using COSMIC (v94) and OncoKB databases [89]. Using a custom code, the homologous protein-coding locations for canine variants were generated for the cancer genes, essentially lifting over the variant position from canine to human. The homologous human variants were analyzed using Functional Analysis through Hidden Markov Models (FATHMM) to predict cancer driver or passenger mutations [90]. The criteria for selecting genes to generate the oncoplot were as follows:

a. Cancer-related genes with recurrent mutations in at least 10% of the samples.
b. Genes with mutations present in only one sample, provided the mutation was predicted to be a cancer driver by FATHMM.
c. Thyroid-related genes with somatic mutations.

The mutational signatures of the samples were explored using the MutationalPatterns R package [91]. The microsatellite instability (MSI) score for each tumor was calculated using MSIsensor-pro [92].

#### Variant validation using Sanger sequencing

Validation of selected variants and gene fusions was conducted using Sanger dideoxy sequencing (Genewiz) of PCR amplified products from genomic DNA or cDNA. Following amplification of identified regions, the amplicons were evaluated by gel electrophoresis, gel isolated, and sequenced using either the forward or reverse amplification primers. Primers were designed from STAR-fusion predicted cDNA sequences of fusion genes or identified variants using Geneious or Primer3Plus (**S2 Appendix**) and purchased from Integrated DNA Technologies (Coralville, IA, USA). Results were aligned with the reference sequence using Geneious (Boston, MA, USA) software to identify variants.

#### Statistical analyses

Statistical analyses were done using R statistical tool (v3.6.1 and v4.1.1) and GraphPad Prism software (v10.4.1). Correlations were assessed by calculating Pearson and Spearman correlation coefficients, and the FDRtool R package was used to correct for multiple testing. The oncoplots and heatmaps were plotted using ComplexHeatmap R package [93]. Bar charts and boxplots were plotted using R package ggplot2 [94]. Clinicopathological data comparisons between FTC and MTC were assessed via nonparametric comparison with a Mann-Whitney U test. Survival analyses (PFI and ST) were performed using Kaplan-Meier curves with log-rank test and COXPH analyses; animals with metastasis noted at diagnosis were excluded from PFI analyses. Significance was set at *p* < 0.05.

#### Conclusions

This study demonstrates notable differences in the gene-expression and mutational landscapes of human and canine TC. While canine MTC may be similarly driven by RET signaling (although, unlike humans, no driver mutations nor fusions were detected in our dataset), canine FTC do not appear to rely on RAS/RAF signaling and display a heterogenous mutational landscape, employing a variety of processes for oncogenesis (including ERBB2 and PI3K, as well as dysregulation of epigenomic and DNA repair pathways). Notably, the EGFR/ERBB family displayed differential expression among canine TC, with FTC demonstrating upregulated expression of ERBB2 and MTC upregulated expression of ERBB4. ERBB2 transcriptomic expression correlated with HER2 protein expression in FTC and increased protein expression correlated with decreased PFI, raising the possible utility of HER2 as a diagnostic, prognostic, and therapeutic marker for canine FTC.

Overall, canine TC appears slowly progressive with a median PFI of around 5 years (61 months) and median survival time of 5.2 years (63 months) after diagnosis/surgery in this cohort. While survival metrics comparing tumor origin (i.e., FTC vs MTC) were not statistically different, the low number of MTC in this study may have diminished our ability to detect differences and further studies are needed. Clinical factors associated with diminished outcomes (particularly in FTC) included larger tumor diameter at diagnosis and use of post-operative adjunctive treatment (i.e., chemotherapy, immunotherapy, and/or radiation therapy). Conflicting with previous studies, metastasis at diagnosis, tumor localization (unilateral vs bilateral) and presence of tumor emboli were not associated with worse outcomes.

## Supporting information

Supplementary plots

## Acknowledgments

Dr. Sarah Tan, for assistance in compiling clinical data from medical records. We also thank Thomas Lee for his initial analysis on canine thyroid transcriptome. We are thankful to have access to the Summit Supercomputer, supported by the National Science Foundation (awards ACI-1532235 and ACI-1532236) at the University of Colorado Boulder and Colorado State University. All NGS analyses were run on the Summit Supercomputer.

