## Supplementary plots for "Comparative multi-omics analysis uncovers contrasting molecular profiles of canine and human thyroid carcinomas"

### Table of contents


| Figure # | Title | Page # |
|---|---|---|
| S1 | Representative photomicrographs of HER2 IHC scoring. | 3 |
| S2 | Normalized expression data of selected differentially expressed genes in FTC and MTC samples. | 4 |
| S3 | Heatmap of over-expressed DEGs in MTC vs Normal samples in dogs. | 5 |
| S4 | Whole exome sequence mapping coverage/depth across 29 soft tissue sarcoma tumors and their matched normal samples. | 6 |
| S5 | Normalized expression PIK3 pathway genes. | 7 |
| S6 | Association of mutational burden and progression-free interval | 8 |
| S7 | Mutational signature analysis of canine thyroid carcinomas. | 9 |
| S8 | Boxplot illustrating the distribution of MEN1 gene expression levels. | 10 |
| S9 | Distribution of *TG* (A) and *CALCB* (B) gene expression levels. | 11 |
| S10 | Distribution of selected fusion gene expression across all 30 samples. | 12 |
| S11 | Kaplan-Meier plots for select clinical parameters. | 13 |

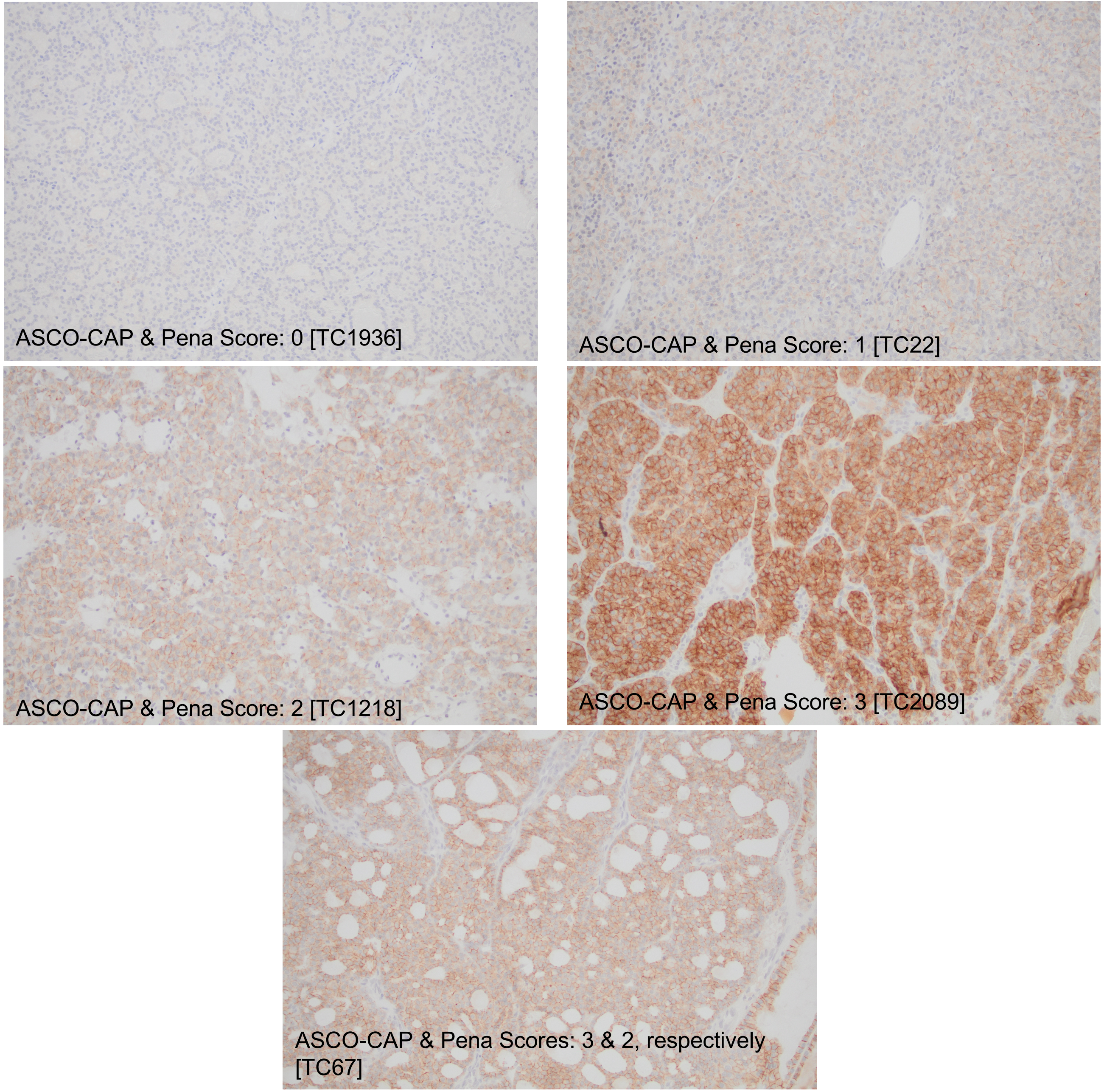


S1 Fig. Representative photomicrographs of HER2 IHC scoring, based on ASCO-CAP and Peña recommendations. 20x objective.

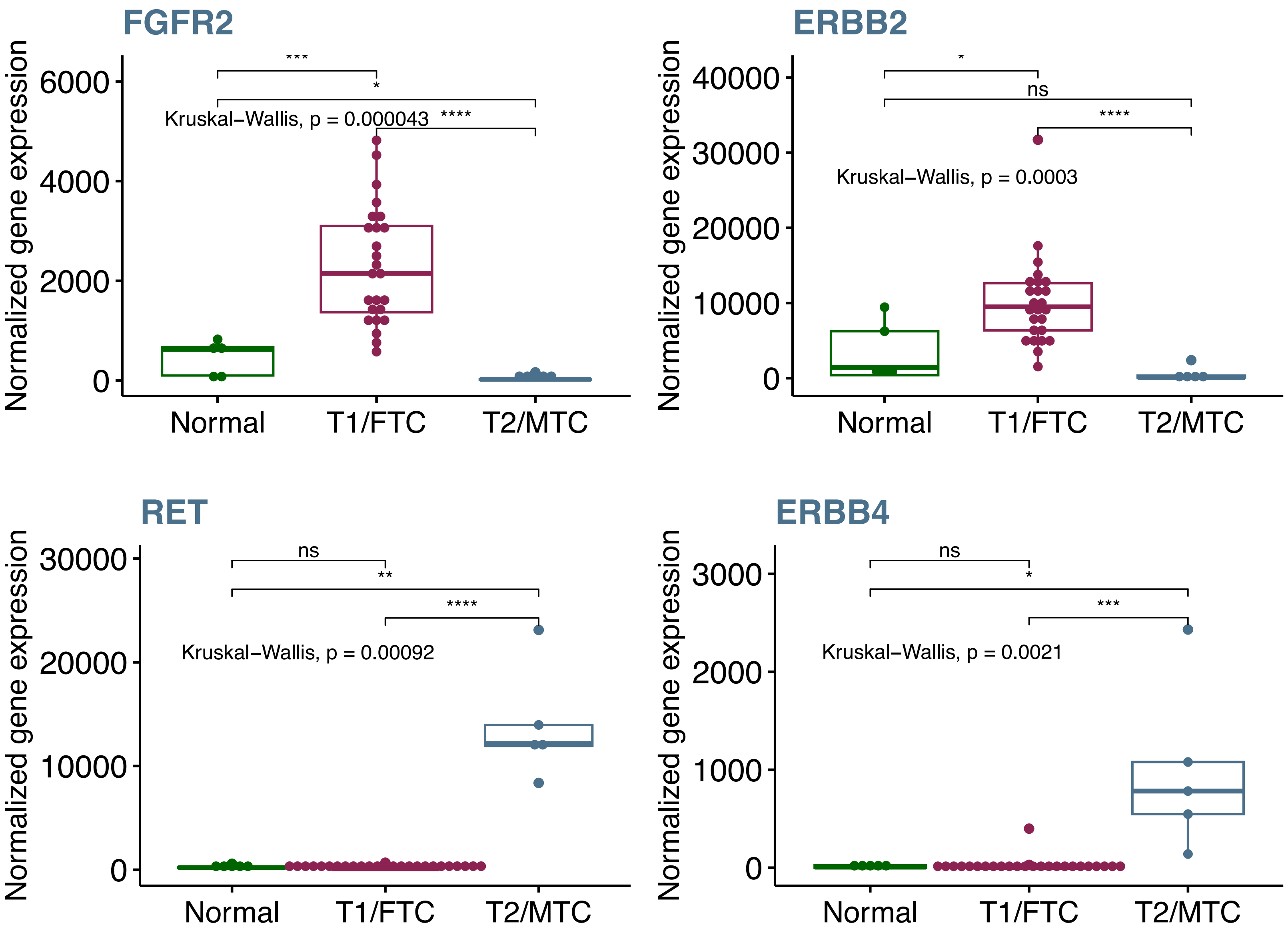


S2 Fig. Normalized expression data of selected differentially expressed genes in FTC and MTC samples. Data from five BarkBase normal samples were included.

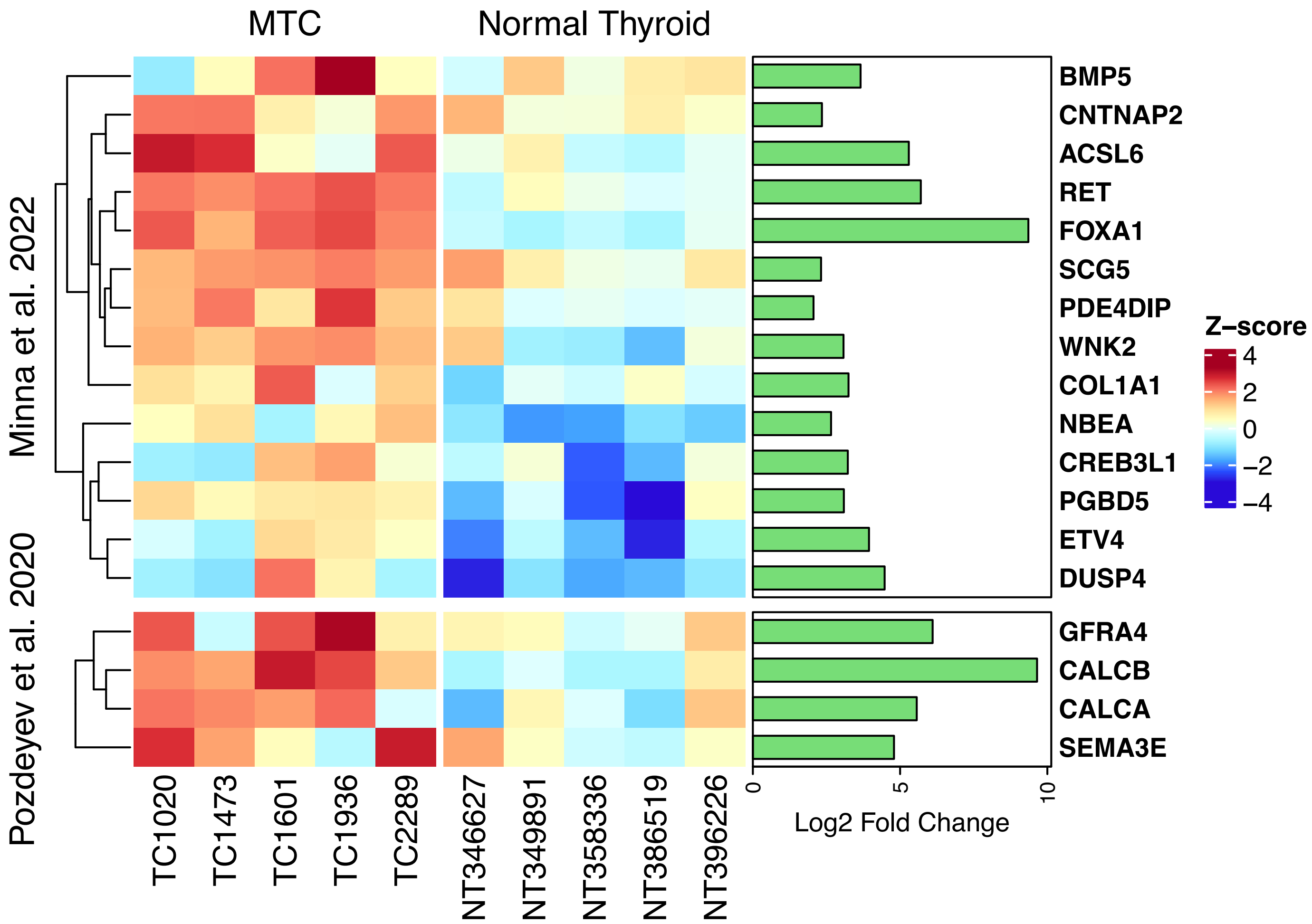


S3 Fig. Heatmap of over-expressed DEGs in MTC vs Normal samples in dogs. The genes plotted here overlap with DEGs identified in human MTC tumors relative to normal thyroid, derived from two published articles. Only cancer-related overlapping genes were plotted in the section labelled Minna et al. 2022.

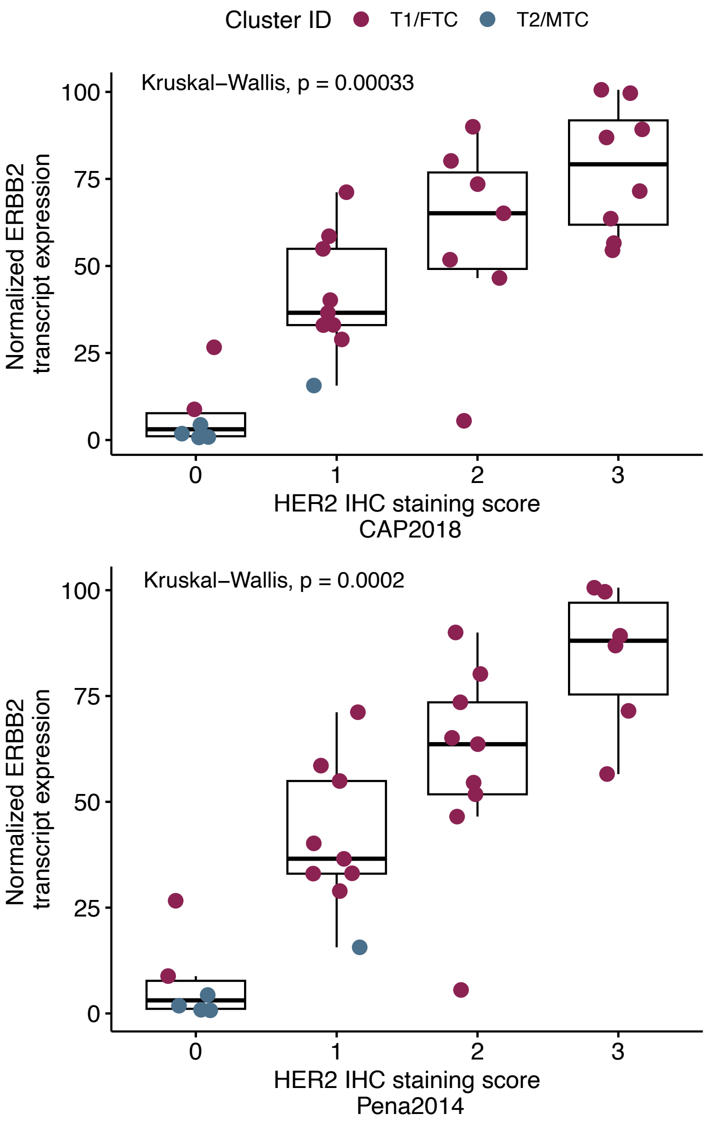


S4 Fig. Boxplot showing distribution of *ERBB2* transcript expression levels across four HER2 IHC scores in canine thyroid tumors.  Two different types of HER2 score were used in this analysis.

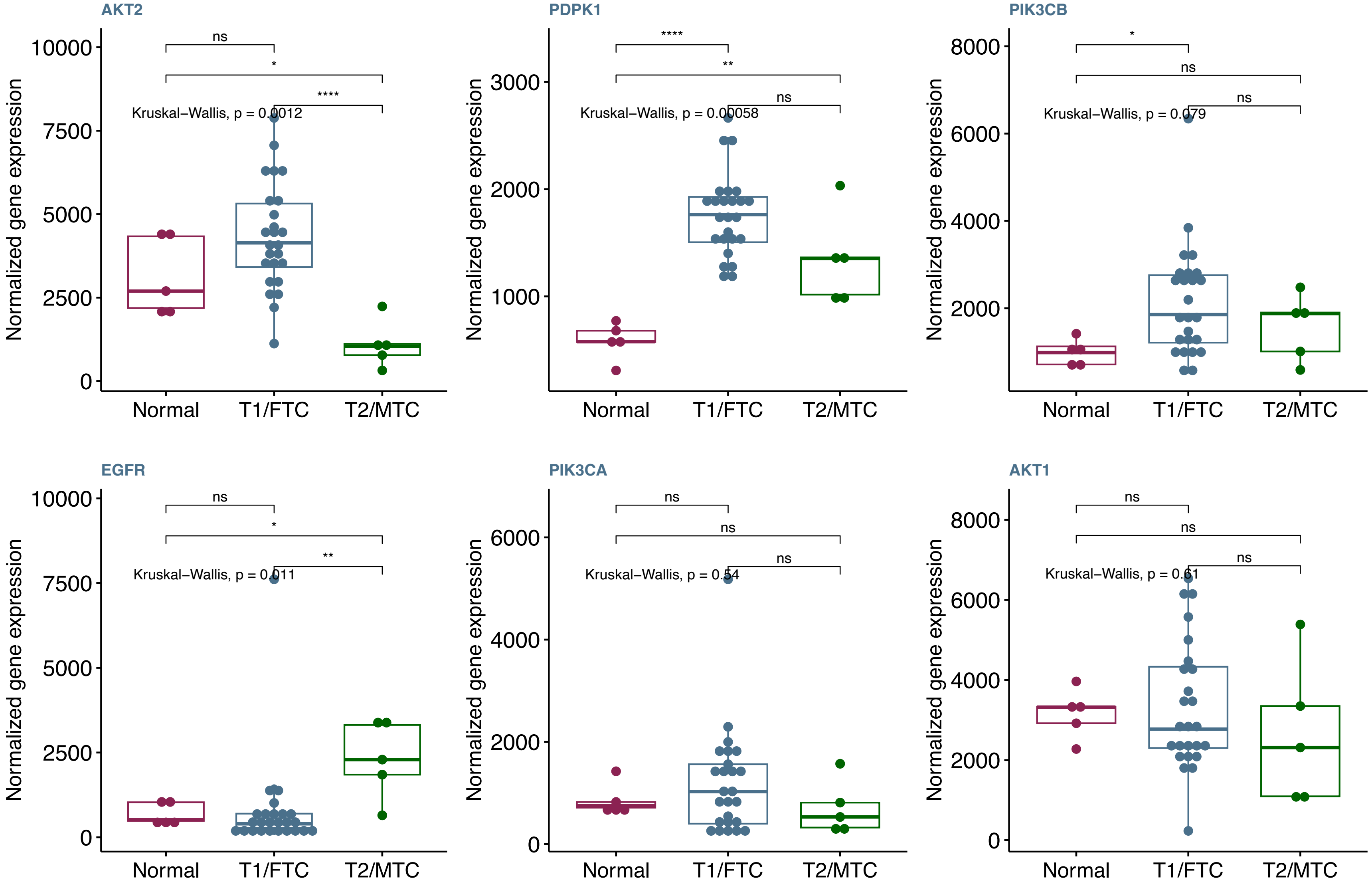


S5 Fig. Normalized expression PIK3 pathway genes. These genes have been previously reported as differentially expressed in canine thyroid tumors.

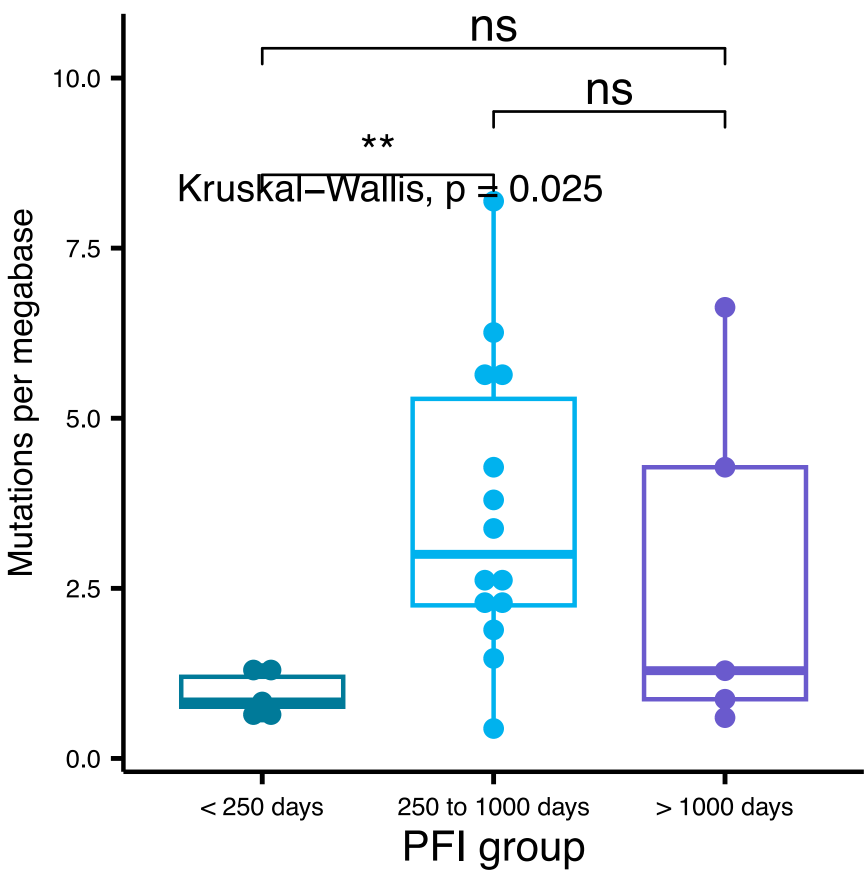


S6 Fig. Association of mutational burden (number of mutation per megabase) and progression-free interval (PFI). Dogs were segregated based on ranges of PFI. Mean number of mutations were highest in dogs with PFI range of 250 to 1000 days.

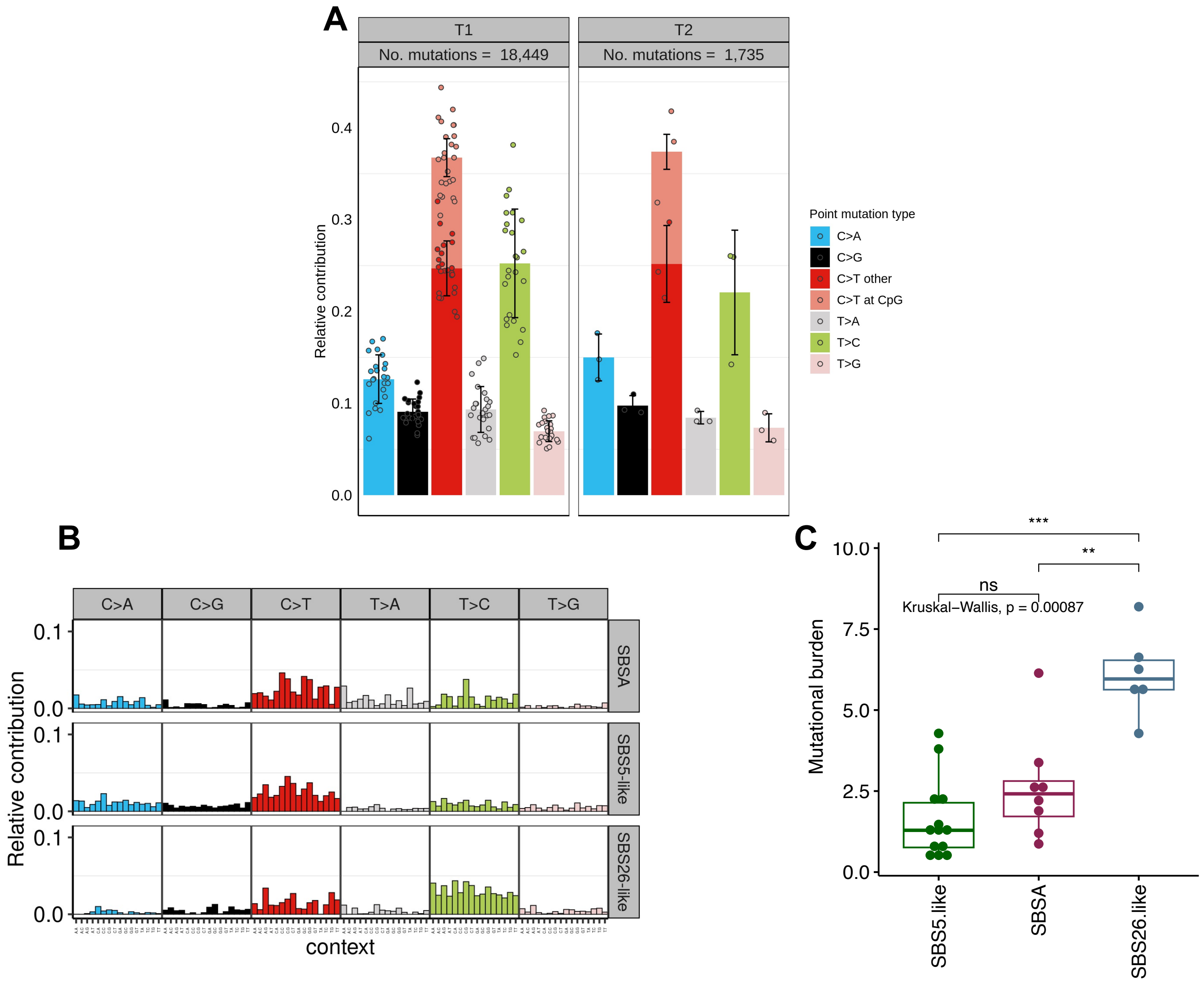


S7 Fig. Mutational signature analysis of canine thyroid carcinomas. A. Relative contribution of 6 single nucleotide substitutions in 25 FTC samples (T1 group) and 5 MTC samples (T2 group). B. Contribution of 96 trinucleotide substitutions to the three mutational signatures identified by NMF analysis of thyroid carcinoma somatic mutation profiles. C. Association of mutational burden to three identified mutational signatures. The dogs with SBS26-like mutational signature had significantly higher mutational burden compared to the other two signatures.

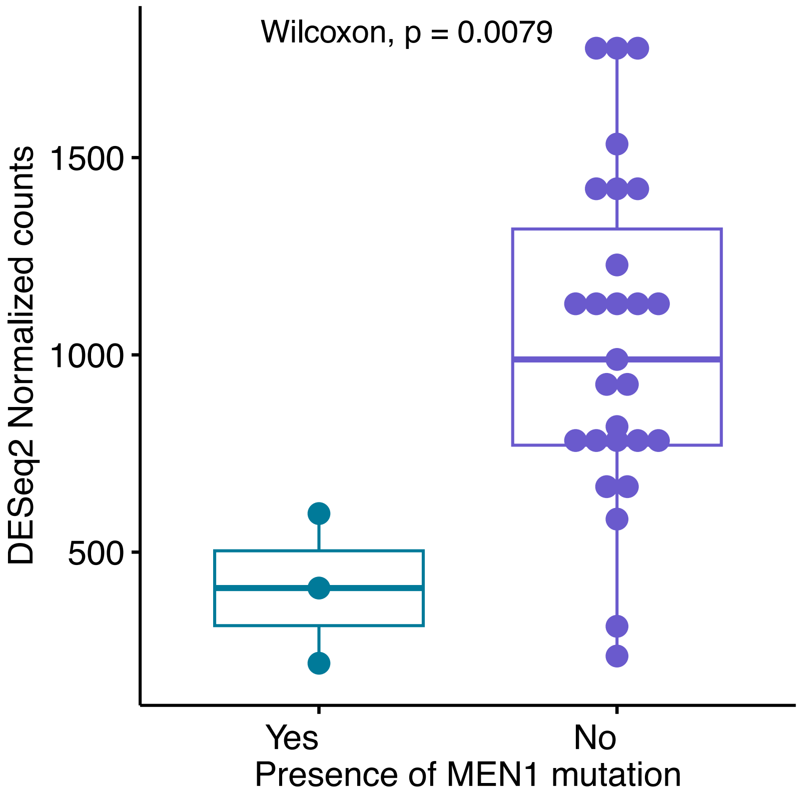


S8 Fig. Boxplot illustrating the distribution of MEN1 gene expression levels. The dogs were grouped by their mutation status (mutant vs wild-type).

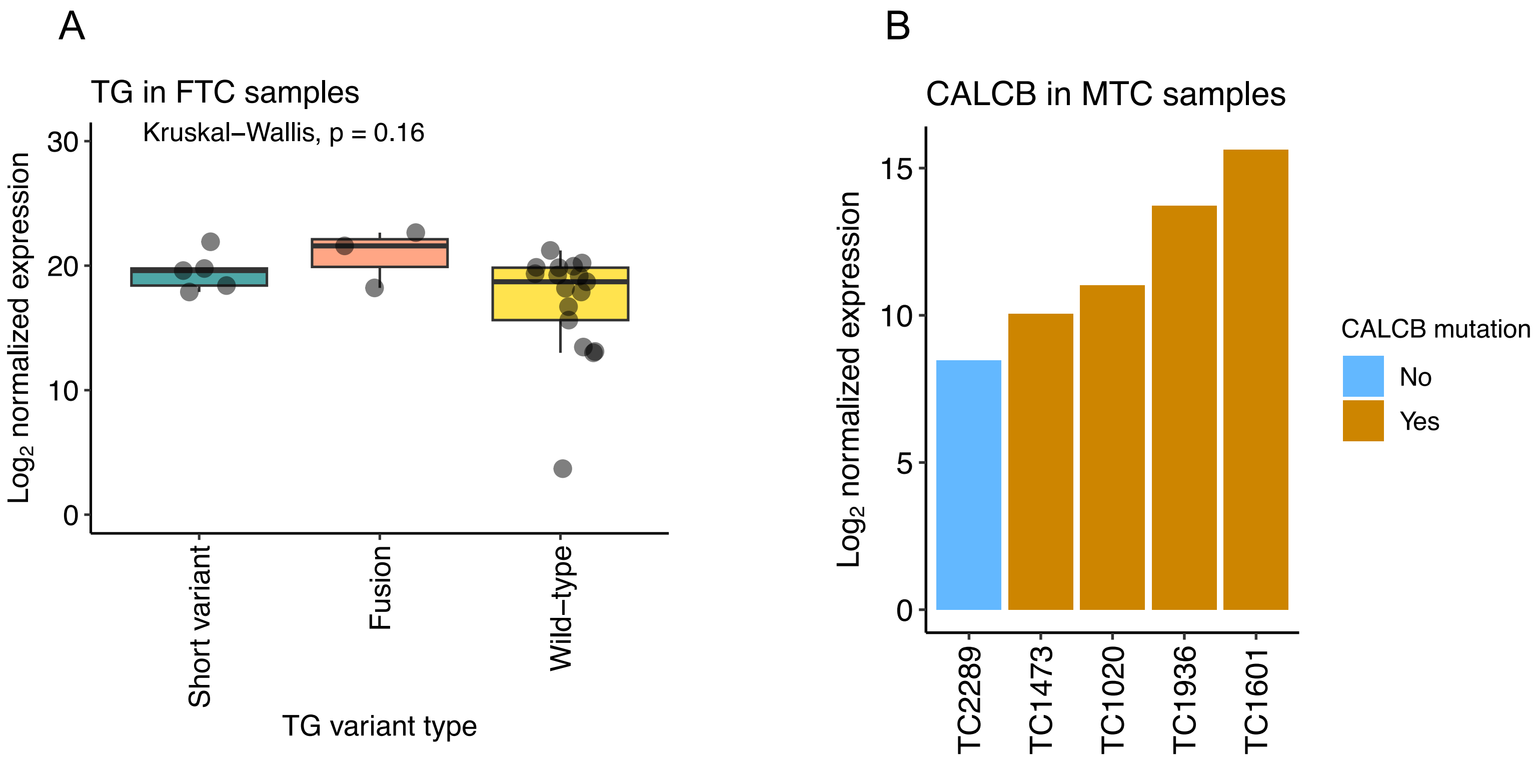


S9 Fig. Distribution of *TG* (A) and *CALCB* (B) gene expression levels. Dogs were grouped by mutation status (mutant vs wild-type) from 25 FTC (A) and 5 MTC (B) samples.

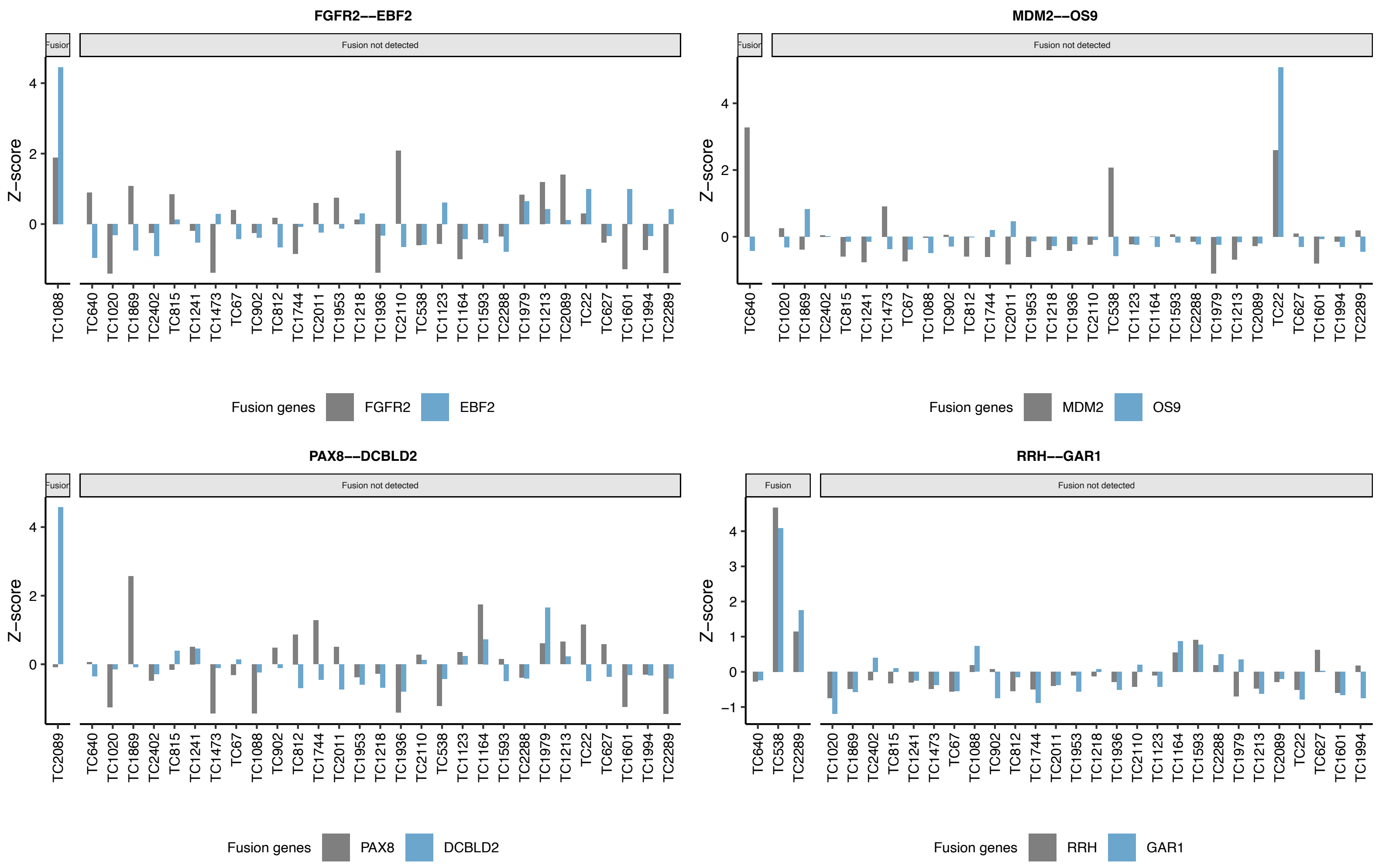


S10 Fig. Distribution of selected fusion gene expression across all 30 samples. These samples were partitioned by the presence or absence of respective fusions.

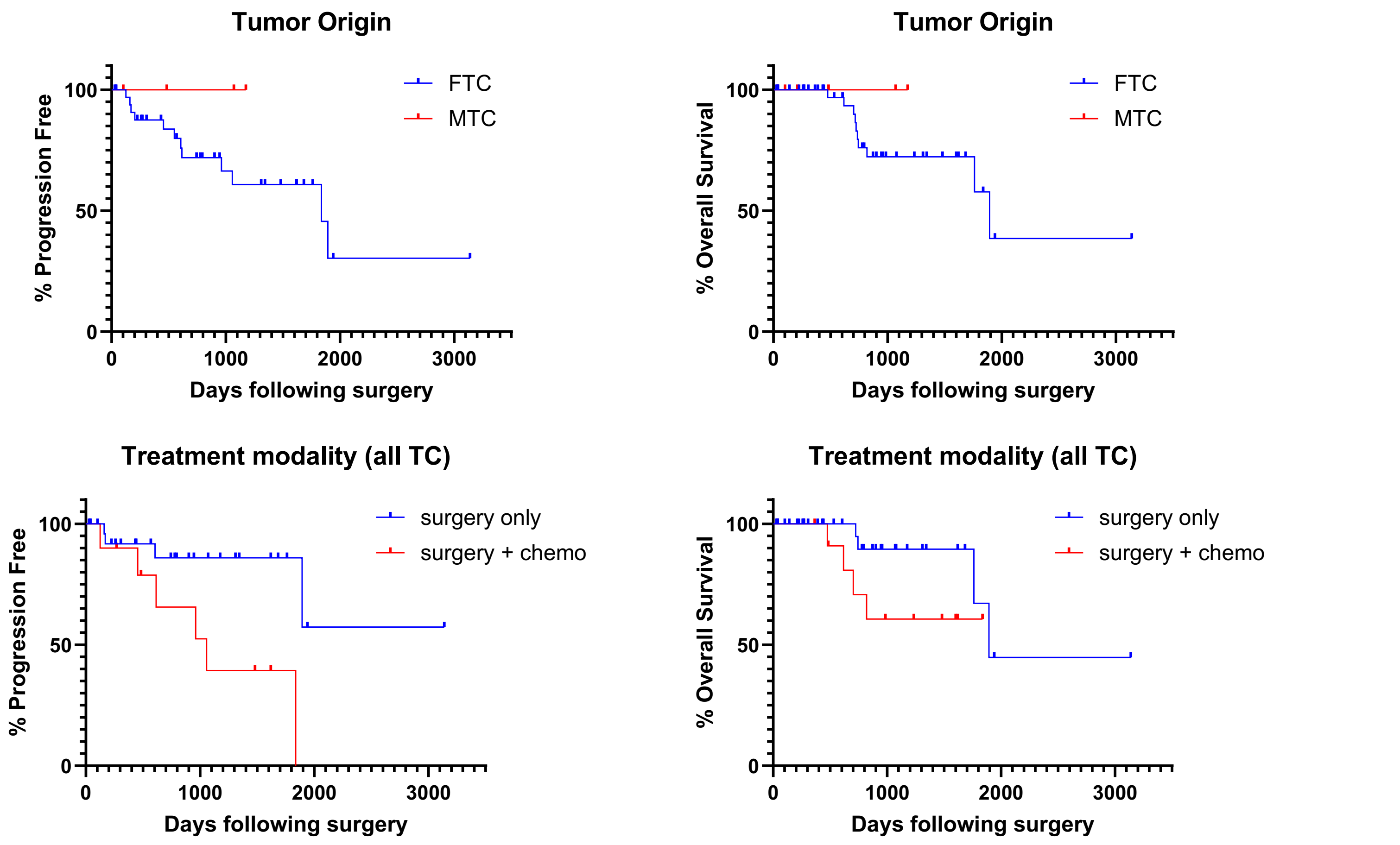


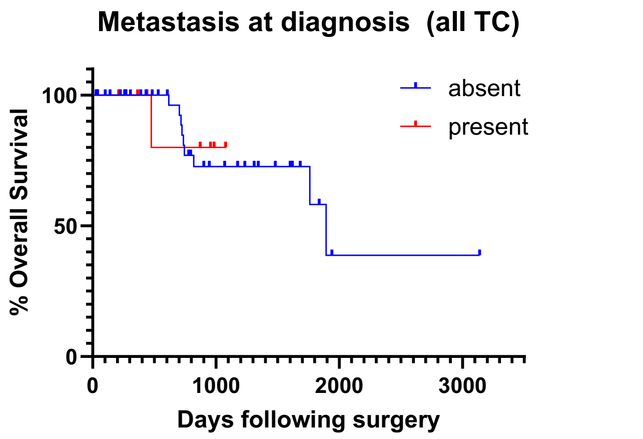


S11 Fig. Kaplan-Meier plots for select clinical parameters. These were not significant on log-rank test ($p > 0.05$).
